# A dynamical systems framework to uncover the drivers of large-scale cortical activity

**DOI:** 10.1101/638718

**Authors:** Arian Ashourvan, Sérgio Pequito, Maxwell Bertolero, Jason Z. Kim, Danielle S. Bassett, Brian Litt

**Affiliations:** Department of Bioengineering, School of Engineering and Applied Science, University of Pennsylvania; Department of Industrial and Systems Engineering, Rensselaer Polytechnic Institute; Penn Center for Neuroengineering and Therapeutics, University of Pennsylvania; Department of Neurology, Hospital of the University of Pennsylvania; Department of Psychiatry, Perelman School of Medicine, University of Pennsylvania; Department of Electrical & Systems Engineering, School of Engineering and Applied Science, University of Pennsylvania; Department of Physics & Astronomy, College of Arts and Sciences, University of Pennsylvania

**Keywords:** dynamical systems, eigenvalue-eigenvector structure, BOLD fMRI, multivariate time series analysis

## Abstract

A fundamental challenge in neuroscience is to uncover the principles governing complex interactions between the brain and its external environment. Over the past few decades, the development of functional neuroimaging techniques and tools from graph theory, network science, and computational neuroscience have markedly expanded opportunities to study the intrinsic organization of brain activity. However, many current computational models are fundamentally limited by little to no explicit assessment of the brain’s interactions with external stimuli. To address this limitation, we propose a simple scheme that jointly estimates the intrinsic organization of brain activity and extrinsic stimuli. Specifically, we adopt a linear dynamical model (intrinsic activity) under unknown exogenous inputs (e.g., sensory stimuli), and jointly estimate the model parameters and exogenous inputs. First, we demonstrate the utility of this scheme by accurately estimating unknown external stimuli in a synthetic example. Next, we examine brain activity at rest and task for 99 subjects from the Human Connectome Project, and find significant task-related changes in the identified system, and task-related increases in the estimated external inputs showing high similarity to known task regressors. Finally, through detailed examination of fluctuations in the spatial distribution of the oscillatory modes of the estimated system during the resting state, we find an apparent non-stationarity in the profile of modes that span several brain regions including the visual and the dorsal attention systems. The results suggest that these brain structures display a time-varying relationship, or alternatively, receive non-stationary exogenous inputs that can lead to apparent system non-stationarities. Together, our embodied model of brain activity provides an avenue to gain deeper insight into the relationship between cortical functional dynamics and their drivers.

## Introduction

Over the past few decades, functional MRI has widened our understanding of the functional organization of intrinsic brain networks and their role in cognition and behavior. Classical univariate (i.e., voxel-wise) analyses of fMRI signal (i.e., blood-oxygenation level-dependent, or BOLD) have been instrumental in probing the specialized function of brain regions. More recent approaches using functional connectivity and network neuroscience portray a complex and multi-scale set of interactions between brain structures. Following this view, a wide array of graph theoretical and complex systems tools have been used to describe BOLD dynamics (e.g., (44; 31; 2)).

Despite these efforts, we still lack a unified mechanistic framework that overcomes three key limitations. First, the features of the BOLD signal that are important for neural activity are unclear. Several prior studies demonstrate a relation between BOLD and slow amplitude features of cortical activity (73; 78; 121), and between BOLD and the hemodynamic response function (HRF) (30; 41). These studies imply that the low frequency component of the BOLD signal contains information relevant to underlying neural dynamics (28; 36), although it is also clear that the signal contains artifact (103; 9). Due to the mixture of signal and artifact in the BOLD time series, it is possible that the common practice of band-pass filtering the BOLD signal at low frequencies may exclude functionally relevant signal (25; 48). Second, many graph theoretic and network analyses are inherently descriptive in nature, and lack the power to give a generative understanding of the relationship between model inputs and outputs (for extensions of these approaches that move beyond description into explanation and prediction, see (6)). Finally, model-based approaches often treat the brain as an isolated system by ignoring external input, or assuming an artificial profile of internal and external noise.

To address these three limitations, we develop a generative framework that explicitly includes exogenous input (e.g., sensory input), and provide evidence that the brain’s activity can be fruitfully understood in the context of its natural drivers. Specifically, we use a multivariate autoregressive model with unknown inputs to capture the spatiotemporal evolution of the BOLD signal driven by external inputs, such as sensory stimuli. These models have been used to characterize and predict the evolution of several synthetic and biological systems (56; 88; 69; 23). Many prior studies use this (54) and similar methods such as Granger causality (47) and dynamic causal modeling (DCM) (44), for understanding the directed functional connectivity of BOLD (107; 91). While some prior studies account for the effect of exogenous input (44; 92), they typically assume a simple known and abstract form of the input function (23). Moreover, the inability of models such as DCM to capture signal variations beyond those caused by the external inputs makes the connectivity estimation highly dependent on the assumed number and form of the inputs (94). In this work, we treat the exogenous inputs to the cortex as *unknown* in the model, and we simultaneously estimate the internal system parameters and unknown excitations leveraging recent developments in linear systems theory (50). To the best of the authors’ knowledge, this is the first use of joint-estimation for an LTI system and its unknown inputs in the context to BOLD dynamics, which allows us to uncover the spatiotemporal structure of the drivers of cortical activity and provides new insights on how the brain responds to the requirements of the ongoing task.

To demonstrate the utility of our approach, we begin with a proof-of-concept where we consider a synthetic example for which we retrieve both the internal system parameters and external inputs to a known noisy LTI system. Next, we test the hypothesis that variations in cortical dynamics during different tasks or cognitive states can be accurately modeled as external excitations on fairly stable interactions between cortical regions. Specifically, we recover the unknown external cortical inputs during rest and task scans for 99 subjects with the lowest motion artifact from the Human Connectome Project (HCP), and find significant task-related changes in estimated inputs showing high spectral similarity to that of known task regressors. Interestingly, we identify task-specific information at higher frequency (> 0.1 Hz) components of the estimated inputs, providing evidence for studying the higher frequency content of the BOLD signal. Finally, we show that different brain systems from our estimated dynamics display diverse and distinguishable profiles of oscillation frequency and dampening.

Lastly, we analyze the important model assumption of system time-invariance over the finite time window used to determine the model parameters. Recently, the nature of non-stationarity of BOLD signal and dynamic functional connectivity has been a topic of scientific debate, as several recent publications paint seemingly contrasting portraits of the stationarity of the processes underlying the brain’s functional dynamics (66; 70; 82; 75; 21; 84). However, to the best of authors’ knowledge, no study has examined the stationarity of BOLD signal in the context of time-varying external inputs and their effects. Here, we propose a novel approach to generate null time series and to test for system stationarity using an LTI model with unknown external inputs. Our simulations demonstrate that even estimations from the dynamics of an ideal LTI system may falsely appear to undergo notable switches in periods where it receives exogenous drivers. Moreover, we study this phenomena in relatively long resting state scans (≈ 14.4 mins) using a sliding window approach (≈ 151 secs), and find that a number of low frequency patterns of activity displayed significantly higher spatial deviations over time across all subjects. Consequently, our observations suggest the presence of a few brain structures displaying significant non-stationarities that cannot be explained by short estimation windows or sampling issues alone. Together, these results provide insight into the drivers of cortical dynamics and factors that contribute to their non-stationarity.

## Results

### Extracting inputs from a synthetic LTI system

We use our method (see details in the Materials and Methods section) to explicitly model the contributions of internal system dynamics and external inputs on the BOLD signal during rest and task. To build an intuition for our method, we begin by estimating the internal system parameters and unknown inputs using data simulated from a synthetic LTI model (Eq. 1) with four states representing four brain regions. We first simulate the dynamics of our model (Fig. 1A) where each region is driven by random noise, and one region is driven by an additional square pulse train (Fig. 1B). Next, we jointly estimate our internal system parameters (4 × 4 matrix of interactions) and unknown inputs (4 functions of time) from the simulations, and accurately recover the frequencies of the pulse train input (Fig. 1C). Further, we see that the estimated system parameters change over time whether or not the inputs are explicitly taken into account, indicating that an open LTI system receiving time-varying external inputs can falsely appear non-stationary when examined at different time periods (Fig. 1D,E). Hence, this approach provides an avenue for understanding the sources of estimation non-stationarities.

**Figure 1.**
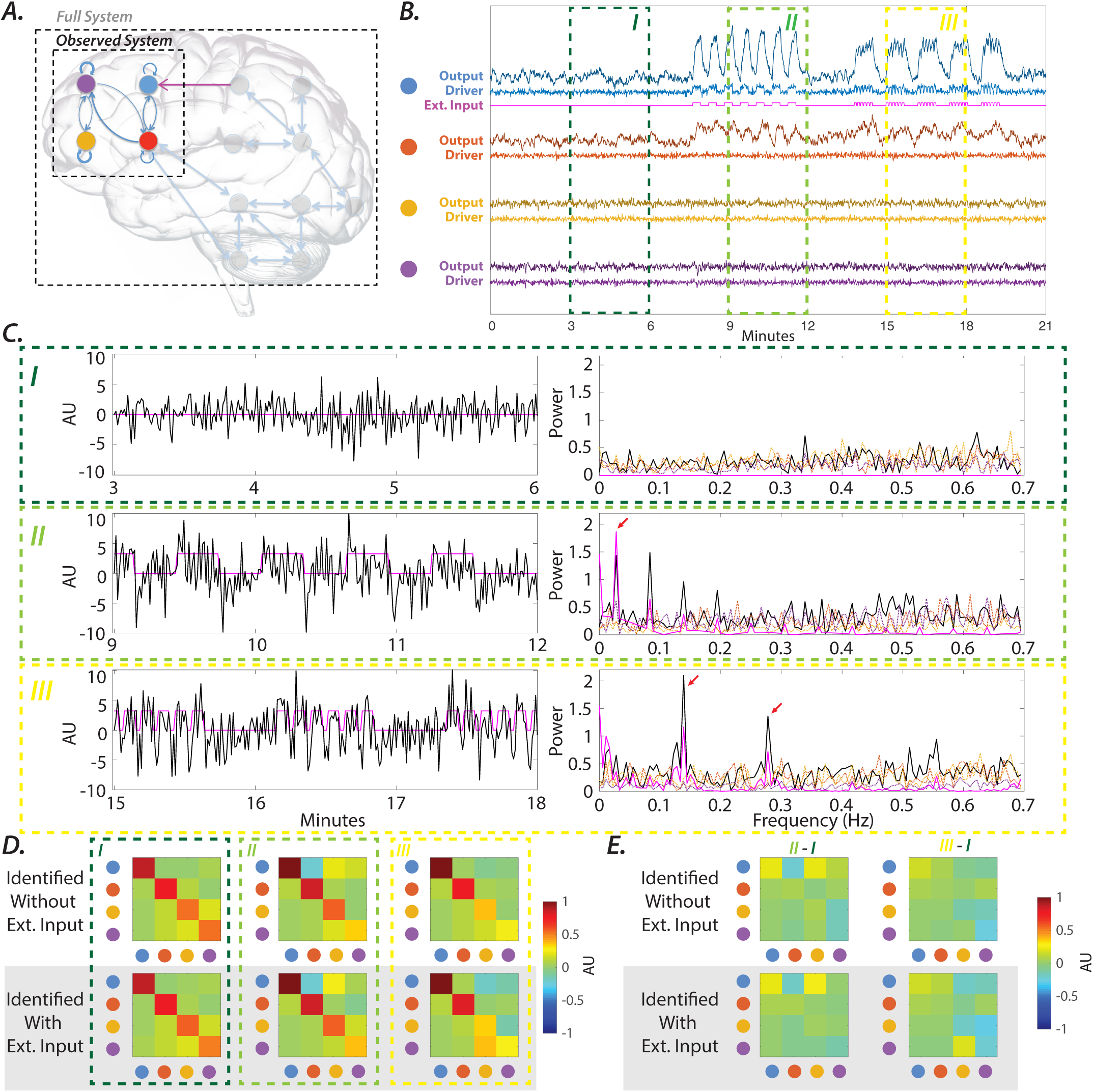
Synthetic LTI system with unknown inputs *(A)* A sketch model of the brain, where we model the activity of only a few regions. The rest of the system including the subcortical structures and peripheral nervous system as well as the external inputs to the brain are assumed the be unknown. Here, we considered a four-dimensional LTI system to describe the dynamics across the four measured sites, whereas the influence of the unobserved system, and indirectly the external stimulus to the full system is assumed to be unknown. The synthetic system was designed with oscillatory modes with 0.01 and 0.06 Hz frequencies (similar to frequencies observed in the BOLD signal), and simulated with 1.4 Hz sampling frequency (similar to the fMRI sampling rate in the HCP dataset). The recorded signal is assumed to be directly reflecting the current state of the system in addition to the random recording noise (SNR = 5). ***(B)*** The color-coded time series (i.e., light blue, light red, light yellow, and light purple) represent the profiles of the nodes’ drivers (i.e., internal noise plus external input), and the darker color-coded time series (time series on top) represent the nodes’ outputs (i.e., simulated system output plus the recording noise). Note that only the blue node receives external input at different points in time (magenta line). Three periods (I–III) are highlighted with color-coded dashed lines. Period I (3–6 mins) is an example of a window with no external stimulations. Period II (9–12 mins) the blue node receives prolonged stimulation (25 samples = 18 seconds blocks, each representing a task condition block), interleaved with similarly sized rest periods (representing inter-stimulus intervals). Period III (15–18 mins) the blue node receives external input, similar to Period II, but with a shorter stimulation window (7 samples = 5.04 seconds) and inter-stimulus intervals (3 samples = 2.16 seconds). ***(C)*** Panels on the left demonstrate the estimated inputs (arbitrary unit (AU)) to the blue node (black line) using our proposed joint-estimation algorithm over three the aforementioned periods. Despite the presence of recording noise and the relatively larger number of input dimensions (m=3) assumed in our model, the estimated inputs capture the trend of the known external inputs to the blue node (magenta line). Panels on the right demonstrate the spectral profile of the external inputs (solid magenta line) and the estimated external input to the blue node (solid black line) and to the other nodes of the system (dashed lines). Note that the low frequency peaks of the estimated external input to the blue node match that of the external input in Periods II and III (red arrows) – an effect that is not observed in other nodes. ***(D)*** Panels from left to right show the system matrix estimated without (top panels) and with (bottom panels) the external input parameter in the model over Periods I, II and III. Recall that the system is fixed across all of these periods, which implies that the changes in the estimated systems are mainly due to the external inputs. ***(E)*** Panels from left to right show the difference between the estimated system matrices over Periods I & II, and I & III, respectively. Interestingly, even the systems estimated using models with the external input parameter similarly failed to capture the true stationarity of the synthetic system.

### Capturing internal and external contributors to BOLD

Now that we have developed an intuition for this method on a synthetic example, we move on to demonstrate the utility of this method for quantifying important spatial and temporal features of both the internal system dynamics and external inputs.

#### Intrinsic brain networks display diverse oscillatory profiles

We begin by showing that the estimated system parameters during resting state reliably capture and reproduce known brain functional organization. Further, because these parameters reside within a quantitative dynamical model, we simultaneously capture both spatial (regions that are co-active) and temporal (oscillation frequency) information through the *eigenmodes* of our estimated system. Specifically, each eigenvector indicates an independent pattern of co-active regions, and its corresponding eigenvalue determines both the frequency of oscillation and change in amplitude of the activation pattern. Intuitively, if we initialize our estimated system state to a pattern of activity corresponding to an eigenvector, then the system states would oscillate and dampen according to the corresponding eigenvalue (see more details in Materials and Methods section).

To capture these patterns of activity, we use our method to estimate the internal system parameters on the resting state time series (1200 TR ≈ 14.5 min) for each of 99 subjects in the HCP dataset. Next, we compute the eigenvectors for all subjects and aggregate these eigenvectors into 10 clusters using *k*-means clustering with *k* = 10 (Fig. 2). Visual inspection of the identified clusters reveals that each cluster consists of one or more canonical resting state networks (RSNs). For instance, the yellow cluster in Fig. 2 contains several regions from the default mode network (DMN), executive control network (ECN), and dorsal attention network (DN). Moreover, each RSN is present in more than one cluster. We quantify these observations by calculating the spatial correlation between the 10 identified clusters and the 7 resting state networks identified in (109), as shown in SI-Fig. 1A. Furthermore, we demonstrate the robustness of the spatial profile of these identified eigenvector clusters across a smaller (*k* = 8, SI-Fig. 1B) and larger (*k* = 12, SI-Fig. 1C) number of clusters. Therefore, these results suggest that the RSNs’ configurations seen in eigenvector clusters capture different functional modes of interactions between RSNs.

**Figure 2.**
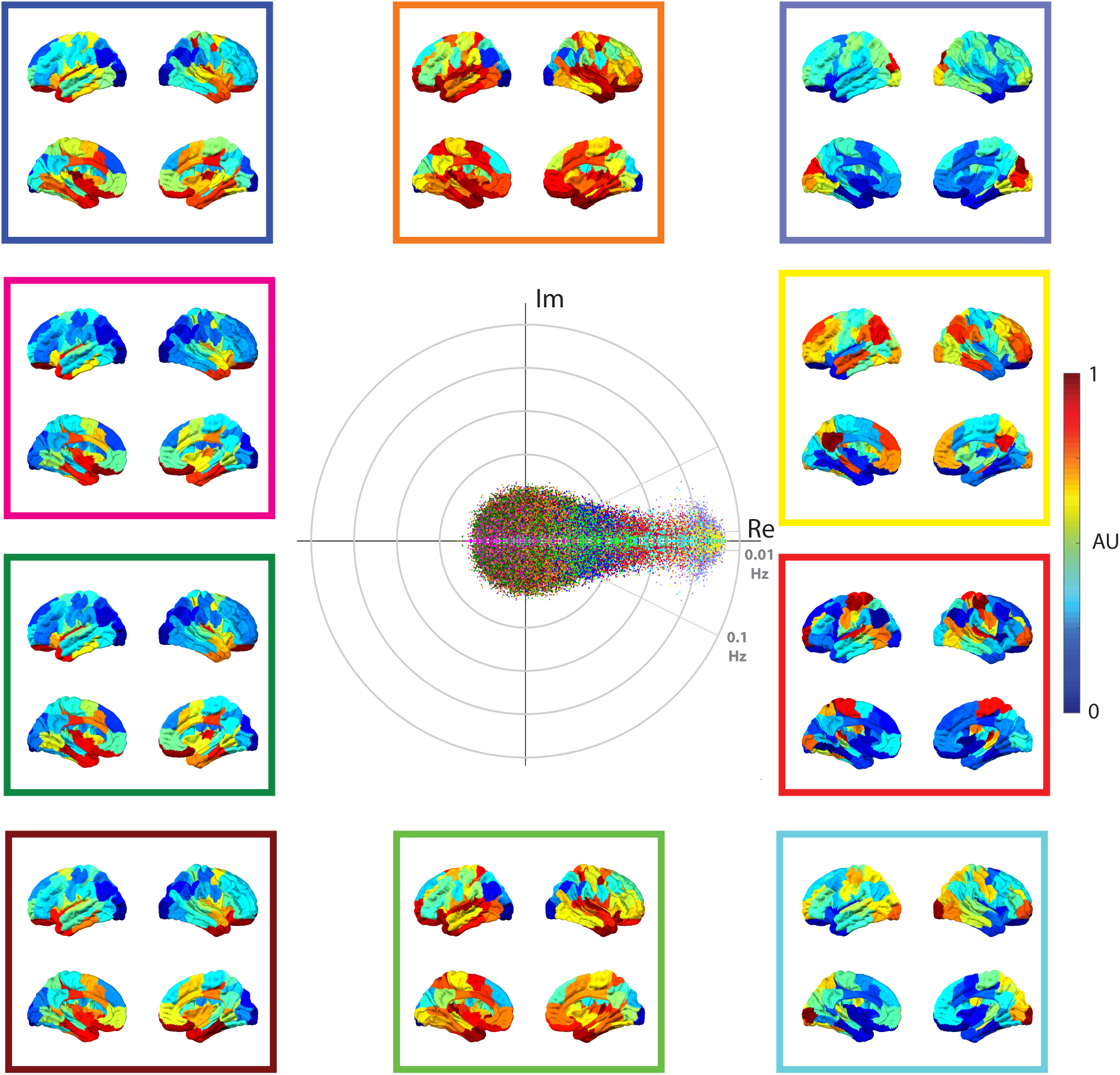
Distribution of eigenvalues estimated from the full (1200 TR ≈ 14.5 min) resting state time series. Clustering the eigenvalues based on their eigenvector’s similarity highlights the spectral profile of different systems. Here, all eigenvectors from all subjects were normalized and clustered into 10 clusters using the *k*-means clustering algorithm. We color-coded the clusters identified across all subjects and sessions (99 subjects × 4 sessions × 100 eigenmodes = 39,600 eigenvalues). The brain overlays represent the spatial distribution of the eigenvector associated with an eigenvalue (displayed with same color code) that is at the centroid of each cluster. To aid the visualization, the centroids were normalized after subtracting each centroid by its minimum element. The color bar represents the normalized values of centroids for all clusters.

Prior research has demonstrated inhomogeneity in the spectral profile of the BOLD signal across the brain (25; 48; 10; 86). As our model also captures temporal features of dynamics, we hypothesize that there is inhomogeneity in the frequency and damping of these clustered eigenvectors. To test this hypothesis, we collect the distribution of eigenvalues corresponding to the eigenvectors in each of the 10 clusters and perform pairwise comparisons (bootstrap *n* = 50, 000, Bonferroni corrected *p* < 0.05). We found a significantly different distribution of eigenvalue frequencies and also damping rates for most cluster pairs (SI-Fig. 2). In addition, the distributions in SI-Fig. 2A reveal that the 10 clusters can be further grouped into three classes based on their frequency and damping rate (i.e., stability). The first group consists of two clusters, one that overlaps with DMN, ECN, and DN (yellow cluster) and the other with Vis (purple cluster), both of which display low oscillating frequencies (average frequency 0.014 ± 0.051 Hz and 0.021 ± 0.065 Hz, respectively), as well as low damping rates (average stability 0.806 ± 0.121 and 0.790 ± 0.127, respectively). The second group consists of two clusters that overlap with SM and DN (red cluster) and with Vis and DN (cyan cluster). These eigenmodes, relative to other clusters, display mid-range damping rates (average stability of red cluster 0.806 ± 0.121 and cyan cluster 0.790 ± 0.127) and frequency distributions with heavy tails skewed towards higher frequencies (average frequency of red cluster 0.103 ± 0.159 and cyan cluster 0.056 ± 0.132). Finally, the third group consists of the 6 remaining clusters, which similarly overlap on the Limbic, VN/Sal, and SM networks (SI-Fig. 1A). These clusters have high average frequencies (average frequencies > 0.14 Hz and standard deviations > 0.15 Hz) and high average damping rates (i.e., low stability with average stabilities < 0.23 and standard deviations > 0.1). Together, beyond classifying these functional clusters of RSNs, these results suggest that the profiles of frequency and damping rate of RSNs’ activity depend on the functional modes of interaction between RSNs.

#### Task scans are marked by task-specific increases in the extra-cortical drivers

In addition to quantifying the spatial and temporal behavior of resting state brain networks, we demonstrate that our method reliably extracts known task regressors from the estimated inputs of each fMRI task (see SI-Fig. 3 for more details regarding the regressors). The external sensory inputs to the brain are believed to be some of the major drivers of cortical dynamics. Therefore, we hypothesize that the inputs to subjects’ brain, as estimated by our method, will mirror real-time changes present in these task regressors.

**Figure 3.**
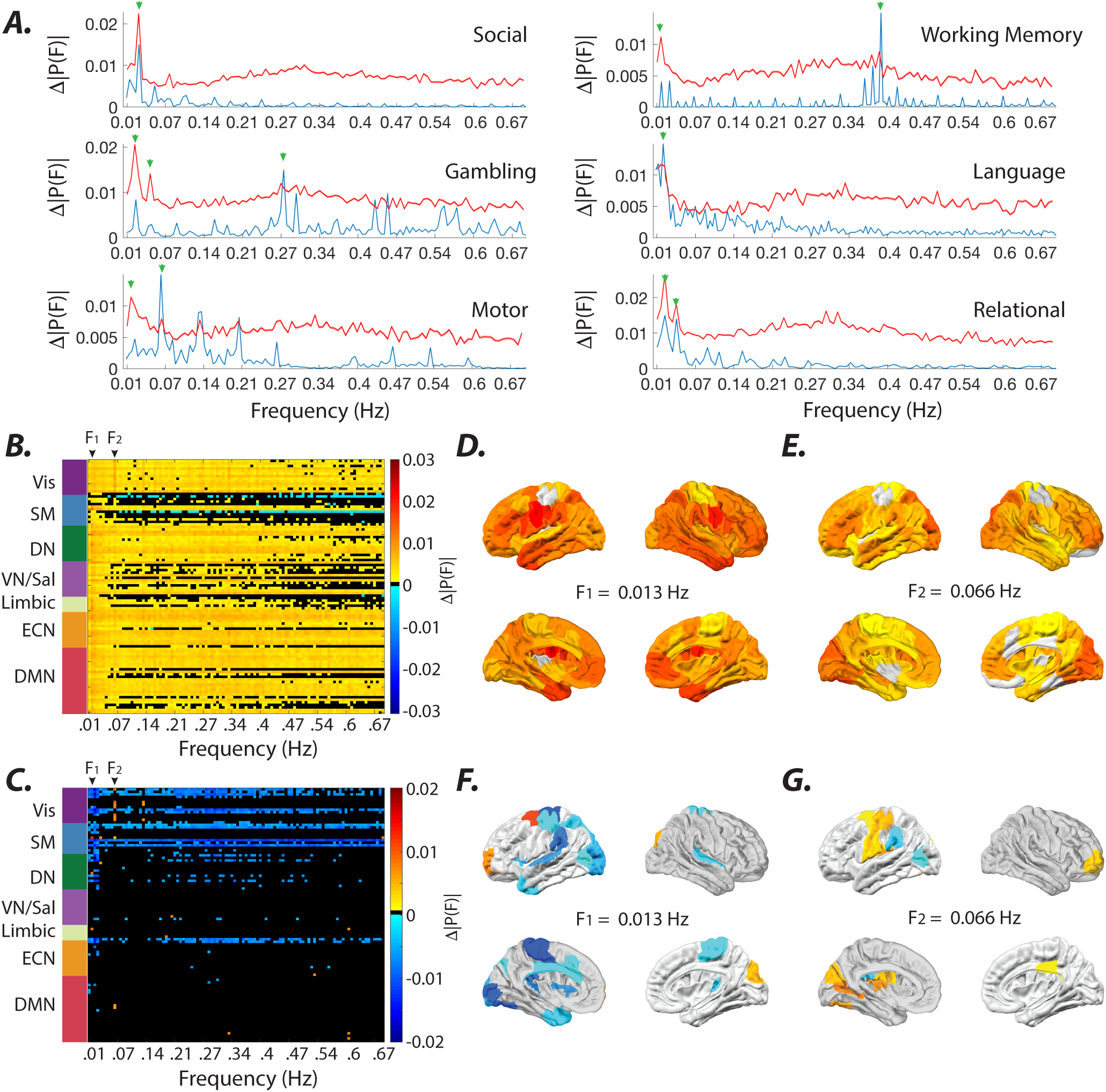
Matching the spectral profile of the known and estimated inputs. *(A)* Red curves show the difference between the average Fourier transform of the estimated inputs to brain regions during the resting state and different task conditions (social, working memory, gambling, language, motor, and relational tasks). Blue curves display the average (two sessions) spectral profile of the known boxcar regressors of each task (see SI-Fig. 3). Note that the peaks of the expected external input frequencies (green arrows) are clearly identifiable across a wide rage of frequencies (even higher than 0.1 Hz). The specificity of these task-related frequency peaks are more clearly highlighted in inter-task comparisons of the estimated inputs, as seen in SI-Fig. 6. ***(B)*** The difference between the average Fourier transform of the estimated inputs to all brain regions during the motor task and resting state. Frequencies for which brain regions did not pass the significance level (Wilcoxon rank sum test, FDR *p* < 0.01) are represented in black. The brain regions are sorted and color-coded (left panel) based on the 7 resting state networks identified in (109), namely the visual (Vis), sensory/motor (SM), dorsal attention (DN), ventral attention/salience (VN/Sal), limbic, executive control (ECN), and default mode network (DMN). ***(C)*** The difference between the average Fourier transform of the estimated inputs to all brain regions during the motor task and other task conditions. Note the significant changes in the power spectrum at expected low frequency and task-specific frequency peaks, such as *F*_1_ = 0.013*Hz*, and *F*_2_ = 0.066 Hz, across several brain regions in both ***B*** and ***C*** panels. Brain overlays in panels ***(D-E)*** and ***(F-G)*** display results in panels ***B*** and ***C*** at *F*_1_ and *F*_2_ frequency peaks, respectively. The colors in panels ***D-E*** and ***F-G*** represent the magnitude of task-related changes in the power using the same color bars in panels ***B*** and ***C***, respectively.

To test this hypothesis, we apply our method to the fMRI activity to estimate the internal system parameters and external inputs for each subject during task performance (social, gambling, motor, working memory, language, and relational) and during rest. Then, we take the difference in the frequency spectra between the average estimated inputs for each task and at rest. We compare this difference to the frequency spectra of the task regressors (Fig. 3A) and find that for each task both the inputs and task regressors share distinctly similar peaks at low (< 0.1 Hz) and high (> 0.1 Hz) frequencies (see SI-Fig. 5 and 6 for the heterogeneity in spatial profile of the external input). Given the significant differences in the estimated inputs between the rest and task conditions, the inputs are unlikely to be driven by inhomogeneity and noise. These findings suggest that the proposed methodology is able to retrieve key temporal properties of the external inputs that serve as drivers of the cortical dynamics.

Beyond the temporal changes in task regressors, we also expect that our estimated inputs will have a larger magnitude of effect during task than at rest. Here we average the absolute value of the estimated inputs across all subjects and sessions for the resting state scans (Fig. 4A), for all task scans (Fig. 4B), and for the difference between the two (Fig. 4C). In general, we see that task conditions are characterized by a significant increase (*t*-test, FDR *p* <0.01) in the average absolute input to the cortex with the exception of several ROIs within the salience network (e.g., the ACC and insula, and hippocampus). This increase is the highest in the visual, attention, and fronto-parietal executive control ROIs. To rule out the recording noise as the source of the estimated spatial profile of inputs, we provided the signal-to-noise map calculated during task and rest in SI-Fig. 4. The dissimilar patterns of average cortical inputs (Fig. 4A-B) and signal-to-noise (SI-Fig. 4A-B) during both rest (Pearson correlation, *r* = − 0.047, *p* = 0.64) and task (Pearson correlation, *r* = − 0.004, *p* = 0.96) provides further evidence of the non-artifactual nature of the estimated inputs.

**Figure 4.**
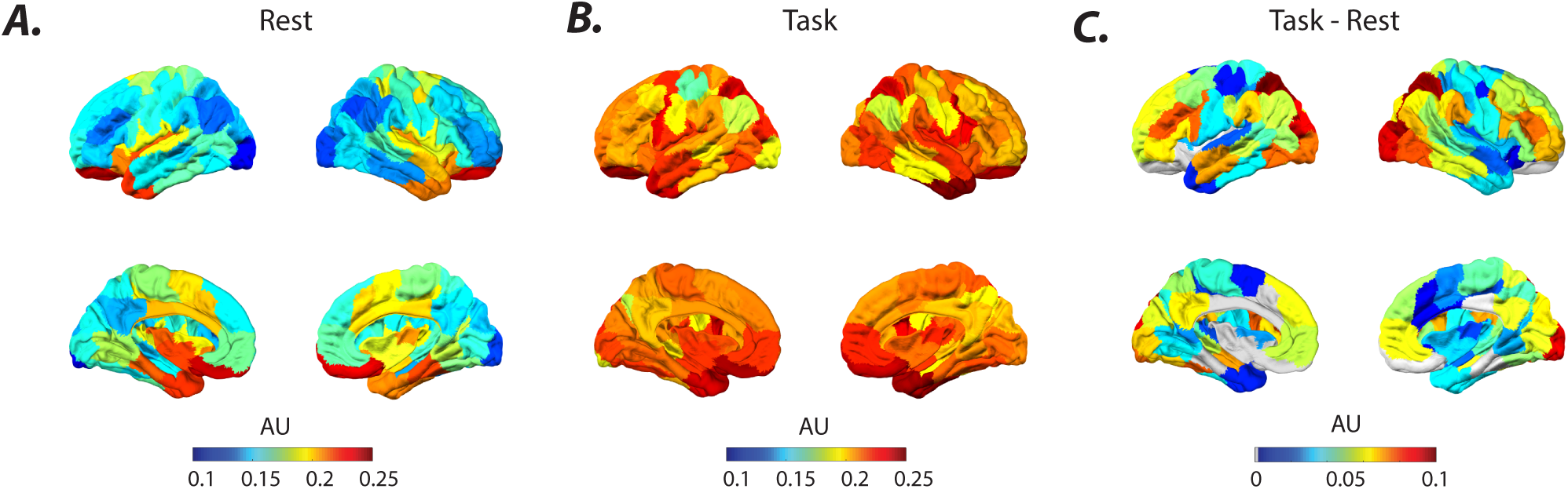
Average cortical excitations or inputs. *(A-B)* The average across all subjects and sessions of the absolute estimated inputs to cortex during resting state and task scans, respectively. ***(C)*** The brain regions with a significant (*t*-test, *p* < 0.01 FDR) increase in the input during task conditions. The regions that did not pass the significance level are colored in gray.

#### Task scans are marked by task-specific changes in the identified system

After capturing both the internal resting state dynamics and the task-based external inputs, we move on to study the internal system parameters estimated from the task-based fMRI. As we demonstrate in our synthetic example, transient inputs to our system can introduce changes in the estimated internal system parameters. Despite these changes, however, we find several consistent and expected differences in the parameters between tasks.

To systematically study these differences, we estimate the internal system parameters and external inputs for each subject and each task condition (including resting state). We first average the system parameters across all subjects to obtain 7 sets of parameters (6 task, 1 rest) and show the change in the average parameters of each task from rest (SI-Fig. 7A). Then, we perform an ROI-wise statistical comparison (Wilcoxon rank sum test, *p* < 0.05 FDR correct for multiple comparisons) between the resting state and task conditions (SI-Fig. 7B) and find significant and expected task-induced changes.

Next, we examine the eigenmodes of our identified system parameters during task fMRI to understand how task-related changes in internal system parameters change system dynamics during tasks. First, we show that the eigenvalues of system parameters identified using a 210 TR window (≈ 2.5 mins) during task conditions display shifts in frequency and damping rate distributions compared to resting state parameters with the same window size. We quantify the changes on the average frequency and damping rate of eigenmode clusters presented in SI-Fig. 12 and show that 9 out of 10 clusters display significant task-related changes (Wilcoxon rank sum, *p* <0.0001). In the same vein, our results highlight a task-related reduction in the number of highly stable (i.e., low damping rate) eigenmodes and an increase in the number of higher frequency (>0.1 Hz) eigenmodes, as seen in SI-Fig. 10. To capture the task-related changes in the spatial profiles of the system’s eigenvectors, similar to Fig. 2, we cluster (*k*-means clustering with *k* = 10) the eigenvectors of the system parameters identified using a 210 TR window during task conditions (SI-Fig. 8) and the resting state (SI-Fig. 9). We find that the spatial profiles of the eigenvector clusters are highly similar between the resting state and task conditions (SI-Fig. 11) given the correlation between cluster centroids, as seen in SI-Fig. 13A. Despite this similarity, statistical comparison (Welch’s *t*-test (119), *p* < 0.05 FDR corrected for multiple comparisons across brain regions) between the spatial profiles of eigenvector clusters reveal task-related changes across all clusters (as seen in SI-Fig. 11). Interestingly, the task-induced changes in the spatial profiles of eigenvector clusters share similarities (SI-Fig. 13B), specifically increased loading on ECN and DMN regions (Fig. 13C). Together, these results highlight the task-related changes in system parameters and demonstrate the spatial inhomogeneity in task-related shifts in the frequency and damping rate of the eigenmodes in these systems.

### Revisiting common processing paradigms and assumptions

Now that we have systematically used this joint estimation method to explore the contributions of internal system parameters and external inputs on the resting state and task BOLD signal, we now use this method to revisit important processing paradigms and assumptions when analyzing BOLD data.

#### Many eigenmodes are present outside of typical filtered frequencies

Commonly, BOLD signals are filtered between 0.01–0.1 Hz because frequencies higher than this band tend to suffer the most from physiological and recording artifacts (for a review on BOLD signal preprocessing see (18)). However, an examination of the eigenmodes from the estimated system parameters for the social task and for the resting state condition (SI-Fig 10) shows many eigenvalues outside of this filtered frequency band (zone A). While several factors such as high frequency recording noise and estimation error due to small window size can contribute to higher frequency (>0.1 Hz) modes, this noise profile is comparable between the resting state and task conditions, allowing us to identify changes in the spectral profile of the system. We find that the task scans are characterized by a significant increase (Wilcoxon rank sum test, p < 0.001) in the average number of eigenmodes with higher frequency (>0.1 Hz) in the medium range decay (0.6 | < *λ* | < 0.8), and a reduction in the number of stable modes with low frequency (<0.1 Hz) with slow decay (0.8 < | *λ* | < 1) across subjects (for details see SI-Fig. 10C). These task-dependent increases in the number of high frequency oscillatory modes highlight the possible functional relevance of these higher frequency (>0.1 Hz) patterns of activation.

#### Non-stationarity of the identified system during rest is spatially heterogeneous

One of the limitations of modeling dynamical processes is the dependence of estimated parameters on the amount of collected data. In LTI systems, estimating system parameters on longer windows of time often yields more accurate slower oscillatory modes, but less accurate fast modes containing crucially important switching dynamics. However, shorter windows result in larger fluctuations of the parameters possibly due to algorithmic instability or the presence of recording noise. As an example, we examine the clustered eigenmodes of the resting state scan (Fig. 2) using shorter windows (210 TR ≈ 2.5 min) and find analogous clusters with similar spatial patterns, as estimated by the full time series (average correlation between matching clusters’ centroids = 0.86 ± 0.2, for details see SI-Fig. 15). The frequency and damping of eigenmode clusters change significantly (Wilcoxon rank sum test, *p* < 0.0001), but depend on the size of the estimation window (SI-Fig. 14). Overall, the eigenmodes of the system identified during the full length resting state time series have lower average frequencies and notably more divergent profiles of damping across all clusters, as seen in SI-Fig. 14. Hence, we cannot determine whether changes in the estimated internal system parameters arise from unaccounted external input (Fig. 1), insufficiently large time windows (SI-Fig. 14), or true changes in the system parameters. To assess the relevance of these various explanations, we compare the fluctuations in the empirical resting state data to those of a true LTI null simulation.

To construct our LTI null, we begin with the empirical resting state time series (1200 TRs) for each subject, and we use our method to estimate the internal system parameters, external inputs, and noise covariance. Then, we use these estimates to simulate a true LTI null time series (1200 TRs). Here, we take these eigenvalues and eigenvectors of the internal system parameters estimated on the full time window (Fig. 5E) to be the most accurate estimate. Next, we characterize the fluctuations in parameters due to shorter time windows by estimating the internal system parameters of short sliding windows (2 TR steps) on the time series for the empirical data, and also the LTI null simulation. For the internal system parameters of each short window, we find the eigenvector in the empirical data (Fig. 5A,C) and LTI null simulation (Fig. 5B,D) that most closely matches each eigenvector of the system parameters of the full 1200 TR estimate (Fig. 5E). We then take the cosine similarity between each eigenvector of the full length window, and all corresponding eigenvectors of the short window estimates, to get a distribution of fluctuations for each full length window eigenvector. SI-Fig. 16 provides a schematic representation of this analysis pipeline.

**Figure 5.**
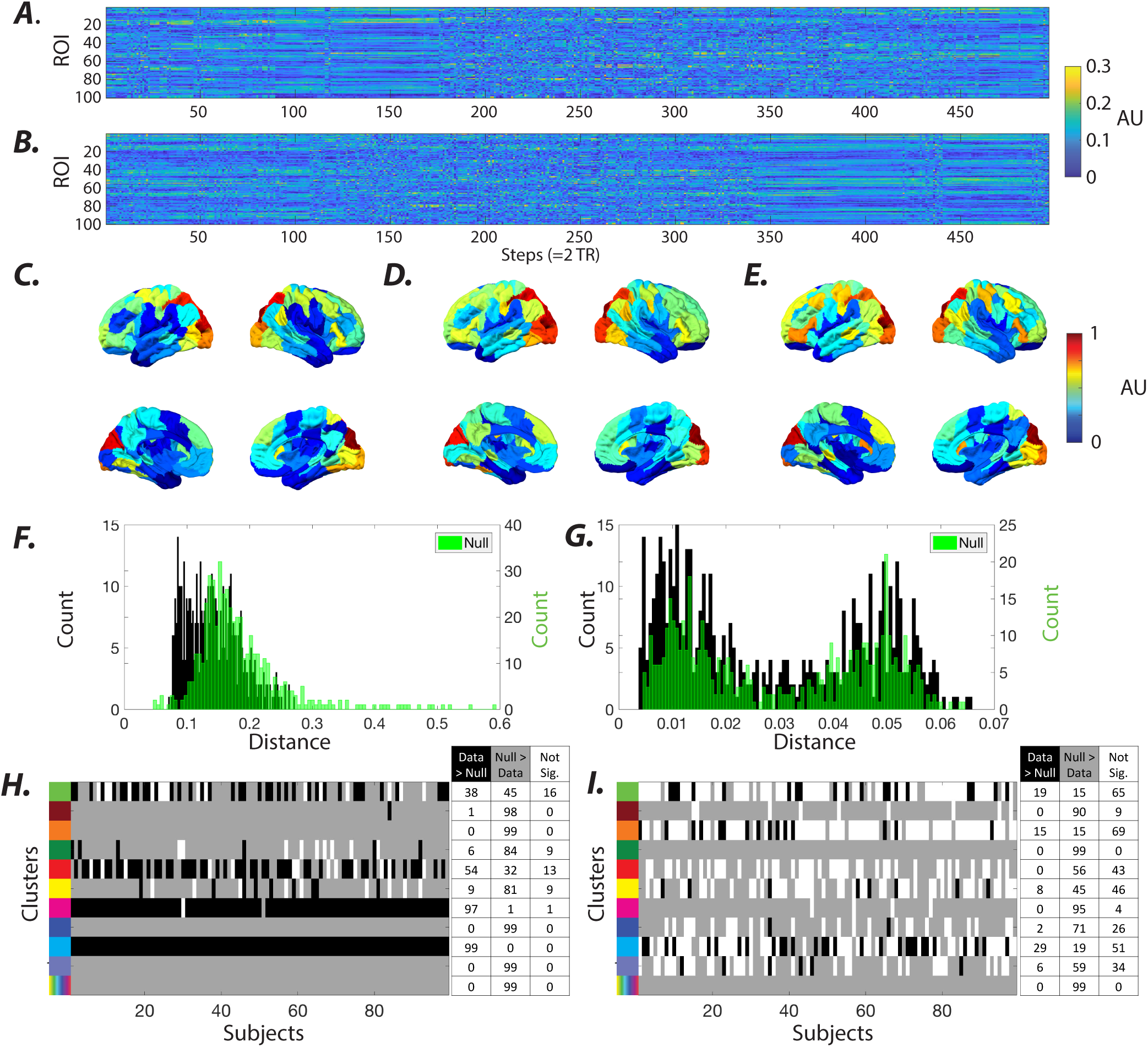
Non-stationarity of the estimated system during rest. *(A)* Sample sorted absolute eigenvectors of the sliding empirical systems, estimated using sliding window (210 TR, with 2 TR increments) from resting state time series. ***(B)*** Sample sorted eigenvectors of the sliding LTI null systems from the same subject estimated from the LTI null time series. ***(C-D)*** The average of the absolute eigenvectors over time in panels **A** and **B**, respectively. ***(E)*** The absolute eigenvector estimated from the full length time series (i.e., LTI null system) with highest similarity to eigenvectors in panels **A** and **B**. To aid the visualization, the brain overlays were normalized after subtracting each by its minimum element. Color bar represents the normalized values of eigenvectors for panels ***C-E. (F)*** Histograms represent the distributions of the distance between the eigenvector in panel **E** and all the eigenvectors in panels **A** and **B**, color-coded in black and green, respectively. ***(G)*** Histograms represent the distribution of the distance between the two adjacent time-points’ eigenvectors for both original data and null time series, color-coded in black and green, respectively. ***(H)*** Non-parametric statistical comparison (bootstrap *n*=50,000, *p* < 0.05, Bonferroni corrected for multiple comparisons across subjects) between the distributions of the eigenvector fluctuations of the sliding empirical and sliding LTI null systems calculated separately for eigenvectors associated with the eigenvector clusters identified in Fig. 2. We measure fluctuations as the distance between eigenvectors of systems identified using sliding window and LTI null systems’ eigenvectors – see SI-Fig. 16 for details). For every subject, rows of eigenvectors with the significantly higher and lower distance to the null LTI systems’ eigenvectors (i.e., fluctuations) are marked by black when the sliding empirical systems’ eigenvector fluctuations are higher than the sliding null (data>null), and they are marked by gray when the opposite (data<null), respectively. Clusters that failed to display any significant difference are marked white. Last row depicts the results for the combined distribution of all eigenvectors for each subject. Note that the vast majority of sliding LTI null eigenvector clusters display fluctuations higher than observed in the empirical systems (especially the magenta and dark green clusters). Nevertheless, the cyan and magenta clusters show the opposite effect in almost all subjects (except two subjects in the magenta cluster). ***(I)*** Non-parametric statistical comparison between the sliding empirical and sliding LTI null’s distributions of the distance (bootstrap *n*=50,000, *p* < 0.05, Bonferroni corrected) between the adjacent sliding-windows’ eigenvectors for all the eigenvectors associated with eigenvalues within the clusters identified in Fig. 2.

Finally, we collect these distributions by eigenvector clusters (according to resting state *k*-means clustering, Fig. 2), and find that overall, there is significantly (bootstrap test, *n*=50,000, *p* < 0.01 FDR corrected for multiple comparisons across subjects) less fluctuation in the empirical data than in the LTI null for the majority of subjects (Fig. 5F,H). However, some eigenvector clusters, including the cyan and magenta clusters actually display significantly (Bootstrap test, *n*=50,000, *p* < 0.01 Bonferroni corrected for multiple comparisons across subjects) more fluctuation in the empirical data than the LTI null across all subjects.

In summary, we find that after explicitly taking into account external input and time window size, we find more empirically measured fluctuation than expected in the corresponding LTI null model in small groups of eigenmodes that spatially overlap the Vis and DN followed by SM and ECN (SI-Fig. 1).

#### Hierarchical organization of inputs to the brain

Because the size of ROIs is a design parameter, it is common to fracture the intrinsically connected networks of the brain into smaller sub-structures using finer parcellations. Therefore, we hypothesize that examination of the correlational structure of the estimated input should reveal that the ROIs within the intrinsically connected networks (or, resting state networks) in general also receive coherent (i.e., correlated) input. Moreover, it has been shown that at the large scale the resting state networks are roughly organized into three functionally coherent groups that form the basis of the main dynamic attractor states of the system (2). Therefore, we conjecture that the coherent output of these large-scale systems are in part the result of the coherent input to these systems.

Examination of the correlational structure of the estimated input during resting state reveals that brain networks receive coherent input (see in SI-Fig. 17B). At larger topological scales all resting state networks (RSNs), except the Limbic network, display significantly higher (Wilcoxon rank sum test, *p* < 0.001) average correlation of inputs between ROIs within each RSN compared to average correlation of inputs between ROIs within and outside RSNs (SI-Fig. 18). Interestingly, some of the RSNs receive anti-correlated inputs on average. Although some of the observed anti-correlation between the ROIs may be artifactual, or the product of the estimation framework, the examination of the estimated inputs during task conditions reveals significant (*t*-test, p < 0.01 FDR corrected for multiple comparisons across ROIs) decrease in the aforementioned global anti-correlation in large number of ROI pairs, see SI-Fig. 17A,C. For example, the average correlation between the estimated inputs to ROIs of SM and Vis RSNs changes significantly (Wilcoxon rank sum test, *p* = 3 × 10^−17^) from low values during resting state (*r* = −0.006±0.157) to higher anti-correlation values during the task conditions (*r* = −0.162±0.192) as seen in SI-Fig. 18B.

Further, the coherent nature of inputs to the brain systems can be inferred from the estimated input matrices – see Materials and Methods for details. In SI-Fig. 19, the principal component analysis of estimated input matrices (concatenated across all subjects and sessions) provides converging evidence of a large-scale topological organization of inputs. Although there are task-related differences between the top three principal components (SI-Fig. 19A-B), resting state and task principal components are very similar as seen in SI-Fig. 19C. Moreover, these principal components similarly consist of one or more RSNs as seen in SI-Fig. 19D-E. For instance, the first principal component calculated from resting state input matrices and the second principal component calculated from task input matrices both overlap mainly with DMN and ECN (SI-Fig. 19D-E). Thus, together, these results support the existence of a hierarchical organization of the input to the brain, where groups of RSNs receive coherent external drivers.

Lastly, we note that correlated residuals in the model may indicate the presence of noise (e.g., autocorrelated recording noise), or the presence of higher-order or non-linear dynamics in the underlying system. In fact, we find that even after accounting for the unknown input, the model’s residuals still contain significant correlational structure (*p* ≈ 0, Li-McLeod portmanteau (LMP) statistic (76)). Regardless, it is interesting to notice that despite the error in the model, we are still able to recover the spectral profile of the task regressors, which further validates the approach. Furthermore, examination of the hierarchical community structure of the adjacency matrix calculated from the correlation between ROIs’ residuals reveal high similarity to the 17 RSNs communities (109), as well as high similarity to hierarchical community structure of ROI input correlations. We quantified the similarity between communities by the z-score of the Rand coefficient (112), as seen in (SI-Fig. 20). Together, these results suggest that a simple first-order linear oscillatory model allow us to capture the evolution of large-scale cortical BOLD dynamics and its drivers, even though more complex models might be able to further explain the internal dynamics of intrinsic brain networks and their task-induced changes.

## Discussion

### The brain is an open system

Based on the theory of embodied cognition, the evolution and the emergent function of the brain can be best understood in the context of the body and its interactions with the environment (111; 26; 89; 19). In this view, the information does not exist in an abstract form outside the agent, instead it is actively created through the agent’s physical interaction with the environment (19). Therefore, understanding the native structure of the external inputs to the brain, as well as the interaction between the brain and its exogenous drivers, is germane to understanding the functional dynamics of the embodied brain (104). From the model identification perspective, we demonstrate that the time varying nature of exogenous (i.e., extra-cortical) inputs can be a contributing source of the apparent non-stationarities in the cortical dynamics. Therefore, our observations provide further evidence that the dynamics of open systems such as the brain should be examined in the context of its drivers.

What are the drivers of BOLD signal? Research shows that cortical output reflects changes in the balance between the strong recurrent local excitation and inhibition connectivity, rather than a feedforward integration of weak subcortical inputs (35). Changes in this balance heavily affects the local metabolic energy demands and consequently the regulation of cerebral blood flow and the BOLD signal, despite the net excitatory or inhibitory output of the circuits (72). Inhibition in principle can lead to both increases and decreases in metabolic demands (for a review on inhibition and brain metabolism see (17)). For instance, inhibition may result in a negative BOLD response when the recurrent excitation is reduced (105; 101; 102), whereas an increase in the local metabolic demands induced by the unaffected input may result in an increase in the BOLD signal (for detailed review see (72)). Moreover, cortical afferents and microcircuits can function as *drivers*, transmitting information about the stimuli, or alternatively as *modulators*, modulating the sensitivity and context-specificity of the response (20; 29). Drivers act through fast ionotropic receptors, whereas modulators originate from various brainstem and basal forebrain nuclei and activate metabotropic receptors with slow and prolonged diffuse postsynaptic effects across cortex (55). Excitatory sensory information, transmitted mostly via glutamatergic or aspartergic drivers, combined with the strong evoked recurrent GABAergic interneurons are a major part of neurotransmission dynamics, which in turn affect the local cerebral blood flow (CBF) (72). Likewise, regulation of cortical excitability mediated by neuromodulatory neurotransmitters including acetylcholine (65), norepinephrine (100; 57; 113), serotonin (65), and dopamine (1; 99) can also significantly effect CBF and the BOLD signal.

Our results reveal a heterogeneous distribution of cortical inputs during the resting state and a significant brain-wide task-related increase in inputs, most pronounced in the attention and executive control system. The results are inline with prior reports of the engagement of these systems in task paradigms, also collectively known as the task-positive networks (39). We also provide evidence that the frequency of transitioning between blocks of task and inter-task intervals are echoed in the spectral profiles of estimated inputs even as high as ≈ 0.2 Hz. The results indicate the sensory and stimulus-related nature of the estimated inputs, yet neuromodulatory inputs may also be a contributing factor. Although neuromodulation may not be stimulus-specific, it can be induced by sensory stimulation (63). For instance, attention strongly effects neural activity and the BOLD signal (27; 61; 13). Despite the nature of cortical inputs, the ability to estimate inputs purely from system outputs by assuming an LTI model of the system is a strong indication that the model can accurately capture the input-output relationship. Still, these results should be interpreted with caution, as the presence of any task-related system non-stationarities may also contribute to the observed task-related frequency peaks in the estimated inputs.

### Large-scale model of the brain as a linear time-invariant dynamical system

Can the brain during resting state scans be fully described as a *linear* and *time-invariant* system? Prior studies demonstrate that temporal fluctuations in the BOLD signal (<0.1 Hz) cannot be fully attributed to linear stochastic processes (123; 45; 83), and suggest that the nonlinearities in the BOLD signal could be attributed to the presence of a strange attractor (45). Examination of the nonlinearity of a *signal* can provide insight into the underlying *system*; however the presence of nonlinearity in a signal does not necessarily imply that the underlying system is also nonlinear. Other neuroimaging studies using paradigms such as “temporal summation” have more directly probed the *system* and provide evidence of *system* nonlinearities (118; 81; 11; 79). To test the linearity assumption, for example, a short and a long stimulus is delivered and the system is deemed linear if the response of the long stimulus can be predicted from the temporal summation of responses of a series of shifted short stimuli. Model-based approaches such as work by (43; 79) have concluded that nonlinear transduction of rCBF to BOLD is sufficient to account for the nonlinear behaviors observed in the BOLD signal. However, care should be take in the interpretation of these results as in the temporal summation framework, where the profile of input is assumed to be known and is approximated by an abstract stimulus representation. Therefore, we believe our proposed joint-estimation framework provides a novel avenue for testing the system linearities through the examination of the estimated unknown inputs in summation paradigms.

Stationary signals are characterized by time-invariant statistical properties, such as mean and variance (24). To date, several tests have been proposed to examine non-stationarity of BOLD time series and the presence of dynamic functional connectivity, including test statistics based on the variance of the FC time series (58; 95), the FC time series’ Fourier transform (53), multivariate kurtosis of time series (66; 70), non-linear test statistics (125), and wavelet-based methods (21; 82), among others (84). These methods commonly test for stationarity by comparing empirically measured properties of the time series to that of a suitable surrogate or null time series that is designed to lack time-varying properties. Null time series are generated by non-parametric resampling (90; 12), phase randomization (67; 53), or simulation of surrogate time series using a generative model of the underlying process, such as vector auto-regression (VAR) approaches (21; 125). Naturally, the choice of null models and measured properties of time series have a profound impact on the outcomes of stationarity tests, which have also contributed to the seemingly conflicting reports on the stationarity of the BOLD signal (82; 75; 66; 70).

Importantly, the presence of non-stationarity in the time series does not directly imply the non-stationarity of the underlying system. The output of an LTI system, for instance, that receives non-stationary external inputs can also display time-varying properties. In this work, we aim to disentangle the non-stationarity of the system from the signal by accounting for the external inputs with unknown profiles as drivers of the system. We extend the commonly used generative autoregressive model and simulated null times series using an LTI model that receives unknown non-stationary external inputs. We test the hypothesis that the human brain during rest is well described as an LTI system with unknown external inputs by comparing the fluctuations of the estimated system parameters using a short (151 seconds) sliding window to that of an ideal LTI null. The ideal LTI null time series was generated with an LTI system that receives unknown external inputs. Parameters of this model including the system parameters, exogenous input, and internal noise were estimated from the full length resting state times series, and residuals of the model were treated as a proxy for the recording noise. Quantification of the expected spatial fluctuations of the sliding window eigenvector estimates of an ideal LTI null time series and non-parametric statistical testing revealed that the majority of the eigenmodes of the system estimated using the sliding window approach during rest displayed spatial fluctuations less than the LTI null. However, a cluster of eigenmodes spanning visual and dorsal attention systems displayed fluctuations significantly higher than the LTI null in all subjects. Our observations suggest that these brain structures likely undergo time-varying changes over the course of resting state recordings. Alternatively, the observed differences can also arise from estimation error, for instance, due to the presence of external inputs, as demonstrated in our synthetic example, or the non-stationarities likely present in the recording noise (34; 87). Although we currently cannot rule out the contribution of the aforementioned factors to the extent of the apparent non-stationarities in resting state (see Methodological considerations), we believe our framework provides a novel avenue for understanding non-stationarities of the BOLD signal based on changes in the estimated underlying LTI system and its external inputs at shorter timescales.

### Spectral profile of brain networks in resting state and task

Historically, a narrow band of slow frequencies between 0.01 to 0.1 Hz was thought to contain information relevant to underlying neural activity, and a higher frequency BOLD signal was considered mainly as artifact (28; 38). More recent evidence, however, portrays a broad band picture of BOLD signal fluctuations with frequencies up to 0.25 Hz (86; 10; 36; 25) and even higher (48). In addition, several studies have characterized the spectral diversity of the intrinsic brain networks (97; 122; 126; 5; 64; 93). The presence of DMN among the slowest oscillating eigenmode during resting state scans with ≈ 0.01 Hz oscillation is inline with the prior reports of DMN spectral profile, as well as the frequencies contributing to functional connectivity of BOLD signals (41; 30). Moreover, we observe a group of eigenmodes with average frequencies around 0.2 Hz mainly overlap with Limbic, Sal/VN, and SM. These results coverage nicely with prior reports of high power (5; 93; 64) and coherence (97) at higher frequencies within these systems. In addition, our approach reveals the gradient in the damping profile of brain system activity (i.e., eigenmodes’ stability), which roughly aligns with the gradient in the spectral profile of these systems.

There are currently a few working hypotheses on what contributes to the documented heterogeneity in the BOLD signals’ spectral profile. For instance, the hemodynamic response function across different brain regions, which is dependent on the diameter and structure of the blood vessels, is heterogenous even across functionally connected regions (14; 52; 80). It has been suggested that the differences in cerebral vascular reactivity (3; 51; 110), baseline venous oxygenation (74), and baseline CBF (68) may all contribute to the spectral differences between RSNs. More recently, alternative theories suggest that RSNs are organized hierarchically according to a sensorimotor-to-transmodal temporal gradient (60; 108; 62). In the same vein, others hypothesize that the differences in the spectral profiles of brain systems may not be merely epiphenomenal, and can be a reflection of the ratio of the excitatory to inhibitory receptors’ densities of the regional populations (114; 93). However, our results demonstrate that RSNs appear in eigenvector clusters with different frequencies, which suggest that RSNs’ activity exhibit different spectral profiles depending on the mode of functional interaction between RSNs. Therefore, if the frequency specificity of RSNs or alternatively RSNs’ large-scale functional modes is a reflection of different aspects of underlying neural activity, future work should aim to understand their structural determinants and link to different rhythms of electrophysiological recordings (16; 77; 22).

Finally, we provide evidence of task-related changes in the estimated system parameters as pointed out by previous studies (e.g. (31)), and further demonstrate that these changes result in significant shifts in the spatial distribution, average frequency, and damping profile of oscillatory modes of the system. Prior research has also reported brain-wide and heterogeneous changes in the BOLD signal power spectrum during task (42; 124; 36). These observations, in theory, suggest that the changes we observe in the estimated system parameters are likely induced by changes in the underlying system and in response to the demands of the ongoing tasks. That said, our simulations also reveal that the mere presence of external inputs can lead to apparent changes in the system parameters in our estimation scheme. Therefore, given that we also document task-specific increases in the profiles of the estimated external inputs to the brain, we attribute the observed non-stationarities partly to the changes in the external inputs, though we do not rule out the possibility of true task-related system non-stationarities. Together, our results suggest that large-scale cortical dynamics, as seen through the BOLD signal, are mostly a reflection of dynamic recruitment of various intrinsic connectivity networks in response to the demands of the ongoing task. Furthermore, the recruitment of intrinsic connectivity networks produces a slow oscillatory and relaxatory signature and entails increased extra-cortical excitations.

### Methodological considerations

We have provided evidence through a synthetic example that the unknown inputs to the LTI system under a random Gaussian noise profile can be estimated with a degree of error depending on the extent of the recording noise. However, structured recording noise such as autocorrelated noise can negatively impact the modeled system (34; 87) as well as the the estimated input. Although, we have included global mean regression (33) as a preprocessing step to account for the common global systemic noise that is present in many of the functional networks (85; 40; 115), our model is unable to account for other unknown structured (e.g., autocorrelated) and non-stationary recording noise (34; 87). Despite these limitations, we demonstrated that our current joint-approximation retrieves valuable information regarding the profile of external inputs, which future work can leverage to shine light on the time-varying properties of the recording noise.

Curiously, we show that the presence of external inputs in our model can still lead to estimation errors that are similar to those observed in LTI models that do not explicitly account for external inputs. Although we leveraged the estimated inputs to better understand the nature of the non-stationarities observed in the BOLD signal, our inability to accurately resolve the parameters of the LTI system under the influence of external inputs is a confounding factor, and critically limits our ability to unravel the sources of nonstationarity. Therefore, the development of system identification tools capable of accurate estimation of system parameters under non-stationarity input and noise conditions is crucial to advancing our understanding of time-varying BOLD signal dynamics. Moreover, our adopted model-based approach does not have the aforementioned limitations of descriptive methods, yet our model is phenomenological, and consequently, describes the recorded sensor dynamics. In addition, we assume that the true functional state of the system is directly observable from the BOLD recording. Future work can extend our proposed framework to account for the latent states of the system based on a biophysical system and observation model (15; 106).

More importantly, since we model the brain as an LTI system, any true transient *switches* or non-stationarities in the system can also appear partially as changes in the inputs to the system. Therefore we can only conjecture whether the observed task-related changes in the system parameters during task scans capture switches in the system that occur rapidly during the short (≈ 30-40 seconds) blocks of task conditions, or slowly over the course of the scanning session, or alternatively, are a reflection of estimation errors due to the presence of external inputs. If the former is true, then we expect some of our estimated input during each condition to also echo the true switches in the system. Future work could focus on assessing the brain during long periods of uninterrupted tasks, such as watching a movie (32; 117), to better understand the nature of the task-related system non-stationarities.

## Conclusions

BOLD signal dynamics can be modeled as a low dimensional LTI system with unknown external inputs during both rest and task periods. We demonstrate that the spectral properties of the estimated parameters of these modeled systems allow us to intuit time-varying behavior of large-scale intrinsic brain systems. Furthermore, we show that our proposed joint-estimation algorithm enables us to better understand the structure of the unknown drivers of BOLD signal fluctuations and shine light on factors that contribute to its apparent non-stationarities. Our results have important implications for how we preprocess fMRI data, but more significantly, our results highlight the importance of studying the brain as an open system. Broadly, our approach provides a framework for understanding the brain’s large-scale functional dynamics and non-stationarities, mechanistically via the modeled system and its time-varying drivers.

## Materials and Methods

### Linear time-invariant (LTI) dynamical systems with external inputs

Each region *i* of interest (ROI) from which the BOLD signal is collected provided us with a time series described by *x*_*i*_[*k*] at sampling point *k* = 0, *…, T*. A total of *n*(= 100) regions are considered and the collection of these signals is captured by the vector *x*[*k*] = [*x*_1_[*k*] *… x*_*n*_[*k*]]^T^, with *k* = 0, *…, T*, which we refer to as the *state of the system* (i.e., it describes the evolution of the BOLD signal across different regions). The evolution of the system’s state is mainly driven by (*i*) the cross-dependencies of the signals in different regions (not necessarily adjacent), and (*ii*) the external inputs that are either excitation noise or inputs arriving from the environment surrounding the regions captured by the state of the system (e.g., stimulus arriving from subcortical structures not accounted for during BOLD signal collection). Subsequently, a first step towards modeling the evolution of the system’s state is:

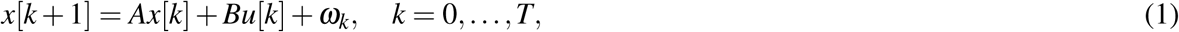

where *A* ∈ ℝ^*n*×*n*^ described the autonomous dynamics, *B* ∈ ℝ^*n*×*p*^ is the input matrix that describes the impact of inputs *u*[*k*] ∈ ℝ^*p*×1^ on the system state’s evolution, and *ω*_*k*_ ∈ ℝ^*n*^ is the internal dynamics noise at sampling point *k*. To restrict the common inputs while minimizing the squared error as well as to empirically capture the inputs to large scale functional networks (Fig. 2), we select a low dimensional input matrix (*p* = 5). Notice that only 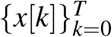 is known since it is the BOLD signal at the different ROIs. In order to determine the *parameters of the system* (1), i.e.,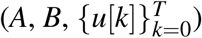, we need to solve an optimization problem that minimizes the distance between the system’s state *x*[*k*] and the estimate of that state given by 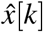 driven by the unknown quantities. Specifically, we have the following optimization problem:

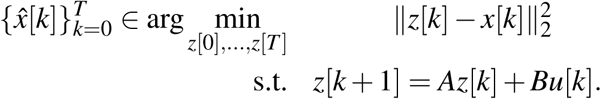

This problem is more challenging than the usual least squares problem considered when the parameters of the system are known (71). Towards solving this problem, we develop a method that, ultimately, boils down to the following steps: (i) we assume that the state *z*[0] = *x*[0], and 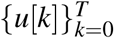 is identically zero, to find an approximation to *A*; (ii) assuming *A* is given by the initial approximation, we provide a structure to matrix *B* and we find an approximation to both *z*[0] and 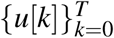, which suffices to obtain *z*[0], *…, z*[*T*] subsequently,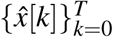; and (iii) assume 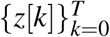 and 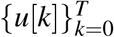 are as approximated in step (ii) and determine an approximation to both *A* and *B*. Therefore, the iterative process consists of executing step (ii) and (iii). Our preliminary tests reveal that the algorithm converges only after few iterations and longer iterations did not provide notable improvement. Therefore, we stop the iterative process after only 5 iterations. To force the inputs to be used as little as possible, since otherwise they could contain the necessary information to obtain the sequence 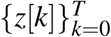 (e.g., consider *A* to be zero and *B* to be the identity matrix), the objective is rather given by 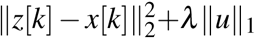, which aims to ensure stationary assumptions measured by the identically distributed errors *ω*_*k*_. The regularization parameter *λ* was obtained experimentally (*λ* = 0.4) while assessing the total mean squared error of the one-step ahead prediction capabilities in a testing subset. Notice that the change of the objective is crucial to ensure that the unknown inputs are close to the real external input to the different ROIs – see more details in (50). Therefore, future work should examine the robustness of our results and test the sensitivity of our observations to different methodological choices, including the dimensions of the input matrix, the size of the system identification window, and the value of the regularization parameter.

### Spectral analysis of an LTI system

Provided an LTI description of the system dynamics (1), the autonomous evolution of the dynamical system can be decomposed in a so-called *eigenmode decomposition*. Briefly, consider the *n* eigenmodes (i.e., eigenvalues and eigenvectors) associated with *A*. Each eigenmode corresponds to an eigenvalue-eigenvector pair (*λ*_*i*_, *v*_*i*_) (i.e., coupled through *Av*_*i*_ = *λ*_*i*_*v*_*i*_), and it describes the oscillatory behavior for a specific direction *v*_*i*_. Specifically, for any given eigenvalue *λ*_*i*_ represented in polar coordenates (*θ*_*i*_, |*λ*_*i*_|), we have that it captures the *frequency* characterized as

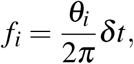

where *θt* corresponds to the sampling frequency, and the *time scale* given by

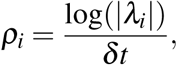

which can be interpreted as the *damping rate*. In particular, we can re-write *A* = *VλV* ^T^, where *V* = [*v*_1_, *…, v*_*n*_] and *λ* = diag(*λ*_1_, *…, λ*_*n*_) are the matrices of eigenvectors and eigenvalues. Subsequently, we can apply a change of variable as *z*[*k*] = *V*^*^*x*[*k*], where *V*^*^ is the transpose conjugate, which implies that 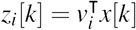 is a weighted combination described by the *i*_*th*_ eigenvector associated with the *i*_*th*_ eigenvalue. Hence, this can be understood as the spatial contributions of the *n* ROIs at a given (spatiotemporal) frequency *f*_*i*_. Additionally, we can revisit the damping rate of the process in such direction *v*_*i*_ by reasoning as follows: first, we can recursively obtain |*z*_*i*_[*k*] | = |*λ*_*i*_| ^*t*^ |*z*_*i*_[0] |. Therefore, we have the following three scenarios: (*i*) |*λ*_*i*_| < 1; (*ii*) |*λ*_*i*_| > 1; and (*iii*) |*λ*_*i*_| = 1. In case (i) and (ii), we can readily see that |*z*_*i*_[*k*] | → 0 and |*z*_*i*_[*k*] | → ∞ as *k* → ∞, respectively. Lastly, in scenario (iii), or practically, when |*λ*_*i*_| ≈ 1, we have that the process oscillates between stability and instability, and we therefore refer to these dynamics as *stable*. In summary, the dynamical process *z*(*k*) describes the spatiotemporal brain BOLD signal evolution. Specifically, the timescales are encoded in the eigenvalues and the spatial contributions of the different ROIs are described by the eigenvectors with a spatiotemporal timescale described by the associated eigenvalues.

### Sorting eigenmodes

Quantifying the fluctuations of the system’s eigenvectors, identified using the short sliding window (210 TR) approach, allows us to highlight the time-invariant features of the system as well as the presence of non-stationarities. For resting state scans, we sorted eigenvectors identified via a short sliding-window based on their similarity to the eigenvectors of the system identified from the full length time series (1200 TR). At every time step, two estimated eigenvectors, one from the short sliding-window and one from the full length time series with smallest distance are paired and then removed. The distance is calculated as one minus the cosine of the angle between the absolute values of the elements of eigenvectors. To sort all pairs of eigenvectors, this process is repeated until all eigenvectors are paired with that of the identified eigenvectors from the full length time series at every time step. Sorted eigenvectors allow us to quantify temporal fluctuations of eigenvectors, and consequently, highlight the time-invariant features of the system. We calculated the distribution of eigenvector fluctuations based on (a) the cosine distance between the sliding-window and full length time series eigenvectors (Fig. 5F), and (b) the cosine distance between eigenvectors of two adjacent time steps (Fig. 5G).

It is worth recalling that, in theory, an LTI system is stationary and thus the identified system should appear identical regardless of the time it was identified. Nevertheless, in practice several factors such as the size of the sampling window, unaccounted external inputs, and recording noise can affect the identification and introduce artifactual fluctuations and non-stationarities even in a purely LTI system. Thus, to better understand the nature of the temporal fluctuations observed in the eigenvectors, we examined and compared the fluctuations of the sorted eigenvectors estimated from simulated LTI null time series to empirically observed fluctuations. The model parameters and their residuals estimated from the BOLD time series were used to simulate the LTI null system’s output and recording noise, respectively. Next, we calculated and compared the distributions of the fluctuations of the null eigenvectors (estimated sliding window approach from the LTI null time series plus recording noise) to that of the empirical data. The empirical and null distributions of the eigenvector fluctuations of all eigenvectors that belong to each cluster identified in Fig. 2 were combined and compared separately for each subject. Bootstrap testing (*n*=50,000, *p* < 0.01 FDR corrected for multiple comparisons across subjects) were performed to establish the statistical significance of all comparisons. This analysis allowed us to identify clusters of eigenvectors that display a high degree of fluctuations compared to the ideal LTI system under comparable external input and recording noise conditions.

### Dataset and Preprocessing

We leveraged data from the Human Connectome Project (HCP). As part of the HCP protocol, subjects underwent two separate resting state scans along with seven task fMRI scans, both of which included two sessions. All data analyzed here came from these scans and was part of the HCP S1200 release. The fMRI protocol (both resting state and task) includes a multi-band factor of 8, spatial resolution of 2 mm isotropic voxels, and a TR of 0.7 s (for more details see (116)). Subjects that completed both resting state scans and all task scans were analyzed. Each of the scanning sessions included both resting state and task fMRI. First, two 15-minute resting state scans (eyes open and fixation on a cross-hair) are acquired, for a total of 1 hour of resting state data over the two-day visit. Second, approximately 30 min of task-fMRI is acquired in each session, including 7 tasks split between the two sessions, for a total of 1 hour of task fMRI (for details see (4)). The 99 subjects with the lowest mean frame wise displacement were used. We utilized a cortical parcellation (*N* = 100 parcels) that maximizes the similarity of functional connectivity within each parcel (98). We preprocessed resting state and task data using similar pipelines. For resting state, the ICA-FIX (96; 49) resting state data provided by the Human Connectome Project were utilized (46), which used ICA to remove nuisance and motion signals. For task data, CompCor (7), with five components from the ventricles and white matter masks, was used to regress out nuisance signals from the time series. In addition, for the task data, the 12 detrended motion estimates provided by the Human Connectome Project were regressed out from the time series. For both task and resting state, the mean global signal was also removed (33).

## Statistics

We performed student’s *t*-test and Welch’s *t*-test (119) to test the statistical significance of the differences between the distributions of interest. Non-parametric Wilcoxon rank sum test (120) as well as the bootstrap (*n* = 50, 000) method (37) where utilized for comparisons of distributions with non-normal profiles. We corrected calculated test statistics for multiple comparisons using false discovery rate (FDR) method (8), as well as the more conservative Bonferroni method (59).

## Acknowledgements

A.A. was supported by The Mirowski Family Foundation. Inc. D.S.B and B.L acknowledge support from the National Institute of Health (R01 NS099348-01) and The Neil and Barbara Smit Fund. D.S.B. also acknowledges support from the John D. and Catherine T. MacArthur Foundation, the Alfred P. Sloan Foundation, the ISI Foundation, the Paul Allen Foundation, the Army Research Laboratory (W911NF-10-2-0022), the Army Research Office (Bassett-W911NF-14-1-0679, Grafton-W911NF-16-1-0474, DCIST-W911NF-17-2-0181), the Office of Naval Research, the National Institute of Mental Health (2-R01-DC-009209-11, R01 - MH112847, R01-MH107235, R21-M MH-106799), the National Institute of Child Health and Human Development (1R01HD086888-01), and the National Science Foundation (BCS-1441502, BCS-1430087, NSF PHY-1554488 and BCS-1631550). Data were provided [in part] by the Human Connectome Project, WU-Minn Consortium (Principal Investigators: David Van Essen and Kamil Ugurbil; 1U54MH091657) funded by the 16 NIH Institutes and Centers that support the NIH Blueprint for Neuroscience Research; and by the McDonnell Center for Systems Neuroscience at Washington University.

## Supplementary Information

**Figure 1.**
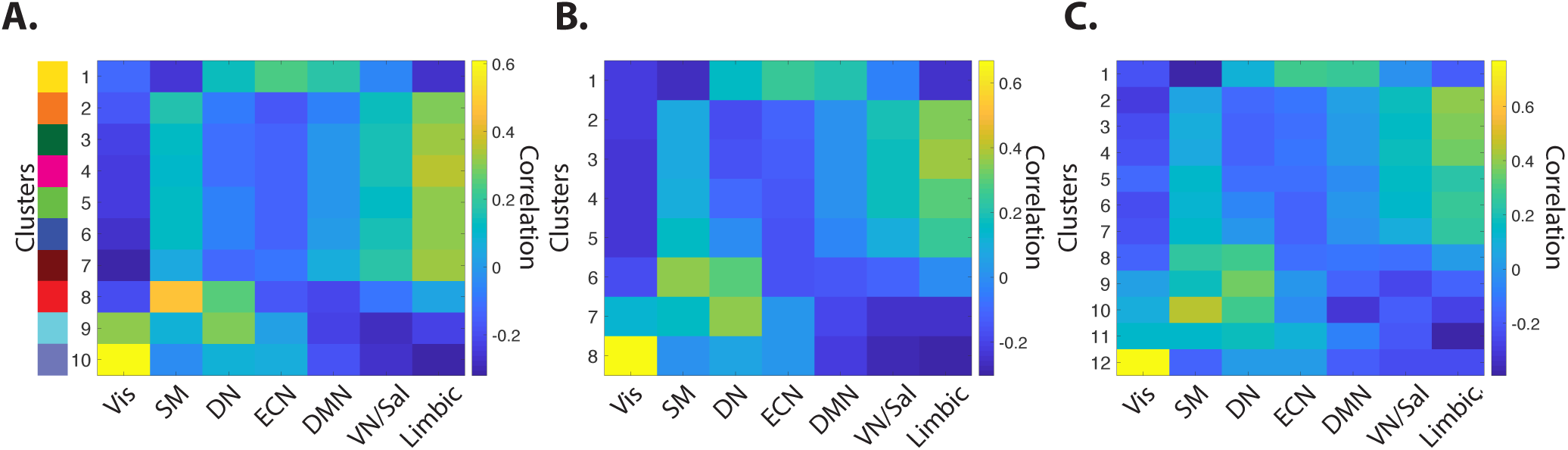
Similarity between eigenvector clusters’ centroids and the resting state networks *(A)* The matrix shows the degree of spatial similarity, which we establish using correlation between centroids of the 10 identified clusters of eigenvectors in Fig. 2 and the 7 resting state networks identified in (109), namely the visual (Vis), sensory/motor (SM), dorsal attention (DN), ventral attention/salience (VN/Sal), limbic, executive control (ECN), and default mode network (DMN). Clusters are color-coded (left) similar to Fig. 2. These results demonstrate that each cluster maps onto two or more resting state networks. Cluster #1 overlaps with ECN, DMN, and DN, clusters # 2–7 similarly overlap with Limbic, VN/Sal, and SM, cluster # 8 overlaps with SM and DN, cluster # 9 Vis, DN, and SM, and finally cluster # 10 mostly overlapping with Vis, followed by ECN, DN, and SM. These clusters are similarly identified at lower (*k* = 8) and higher (*k* = 12) number of clusters as seen in panels ***B*** and ***C***, respectively. Some clusters such as clusters # 1 and # 10 in panel ***A*** are found reliably across different *k* values, though increasing *k* divides other clusters into spatially overlapping subgroups. For instance clusters # 2–5 in panel ***B*** are similarly identified as clusters # 2–7 in panels ***A*** and ***C***. These results demonstrate the robustness of the identified clusters to different *k* values. Moreover, it demonstrate that the eigenvector cluster capture various modes of functional interaction between the RSNs.

**Figure 2.**
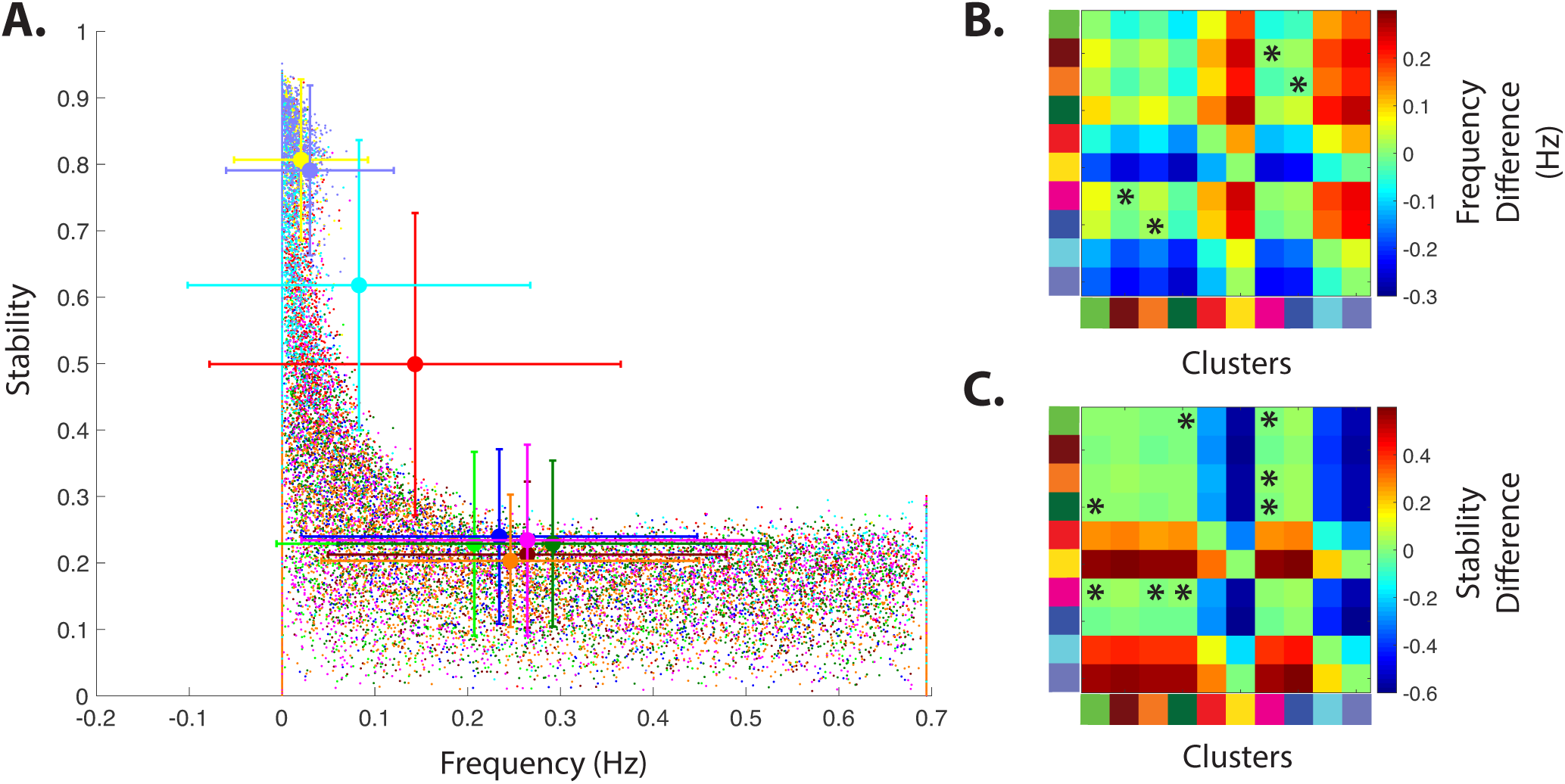
Distribution of frequency versus stability of eigenvalues during resting state *(A)* Distributions of eigenvalues estimated from full resting state time series, based on their frequency and stability (i.e., absolute of eigenvalue). Eigenvalues are color-coded similar to Fig. 2, based on the 10 eigenvector clusters. Error bars represent the mean and the standard deviation of each cluster’s frequency and stability. ***(B)*** Matrix shows the difference between the average frequency of all pairs of clusters. Non-parametric statistical testing (bootstrap *n* = 50, 000 (37), Bonferroni corrected *p* < 0.05) reveal that most clusters, with the exception of few comparisons (highlighted by ‘*’), have significantly different average frequencies. ***(C)*** Matrix shows the difference between the average stability values of all pairs of clusters. Similar to panel ***B***, non-parametric statistical testing (bootstrap *n* = 50, 000, Bonferroni corrected *p* < 0.05) also reveal that most clusters, with the exception of few comparisons (highlighted by ‘*’), have significantly different average stability values.

**Figure 3.**
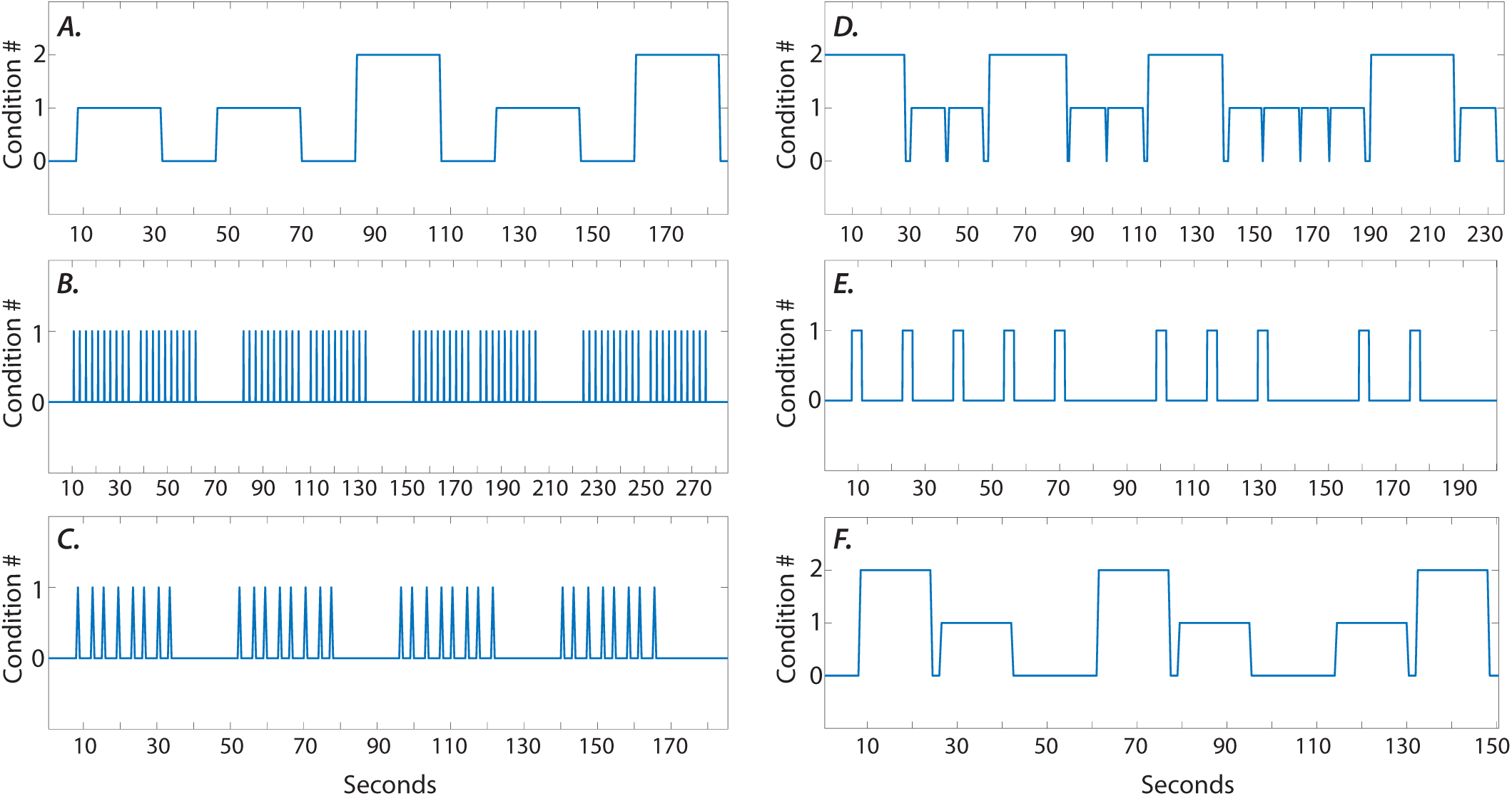
Sample session’s task regressors *(A-F)* A sample scanning session’s boxcar regressor for different task conditions (social, working memory, gambling, language, motor, and relational tasks, respectively). Different task cues and/or blocks are marked by ‘1’ and ‘2’ on y-axis, whereas ‘0’ represents inter-stimuli rest intervals. Since the spectral profile of the task regressors are identical across subjects, we used regressors in **(A-F)** for our analysis.

**Figure 4.**
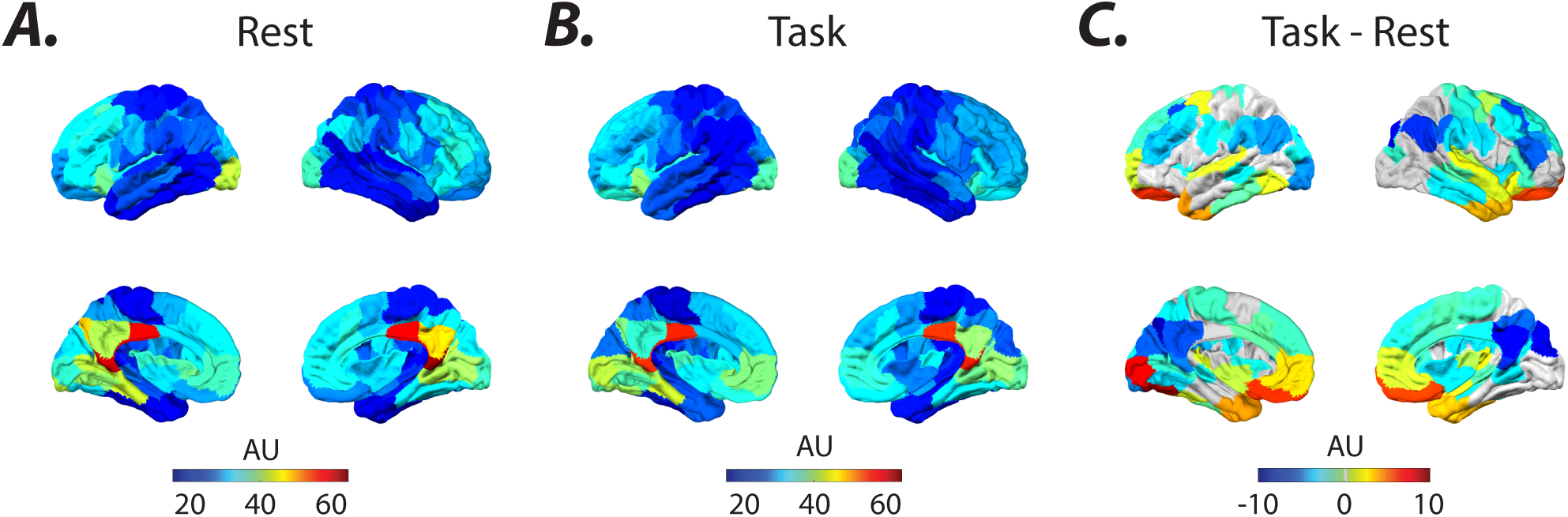
Average signal-to-noise maps *(A-B)* Brain overlays represent the average standard deviation of the preprocessed BOLD signal (prior to normalization) across all subjects and sessions during resting state and task, respectively. The dissimilar spatial distribution of input (Fig. 4) and the signal-to-noise maps in both task and resting state (panels *(A-B)*) suggest that the estimated inputs are not an artifact of the baseline recording noise. ***(C)*** Brain overlay highlights regions that display significant (Welch’s *t*-test, FDR, *p* < 0.01) task-related change in the standard deviation of BOLD signal. Regions that did not pass the significance level are colored in gray.

**Figure 5.**
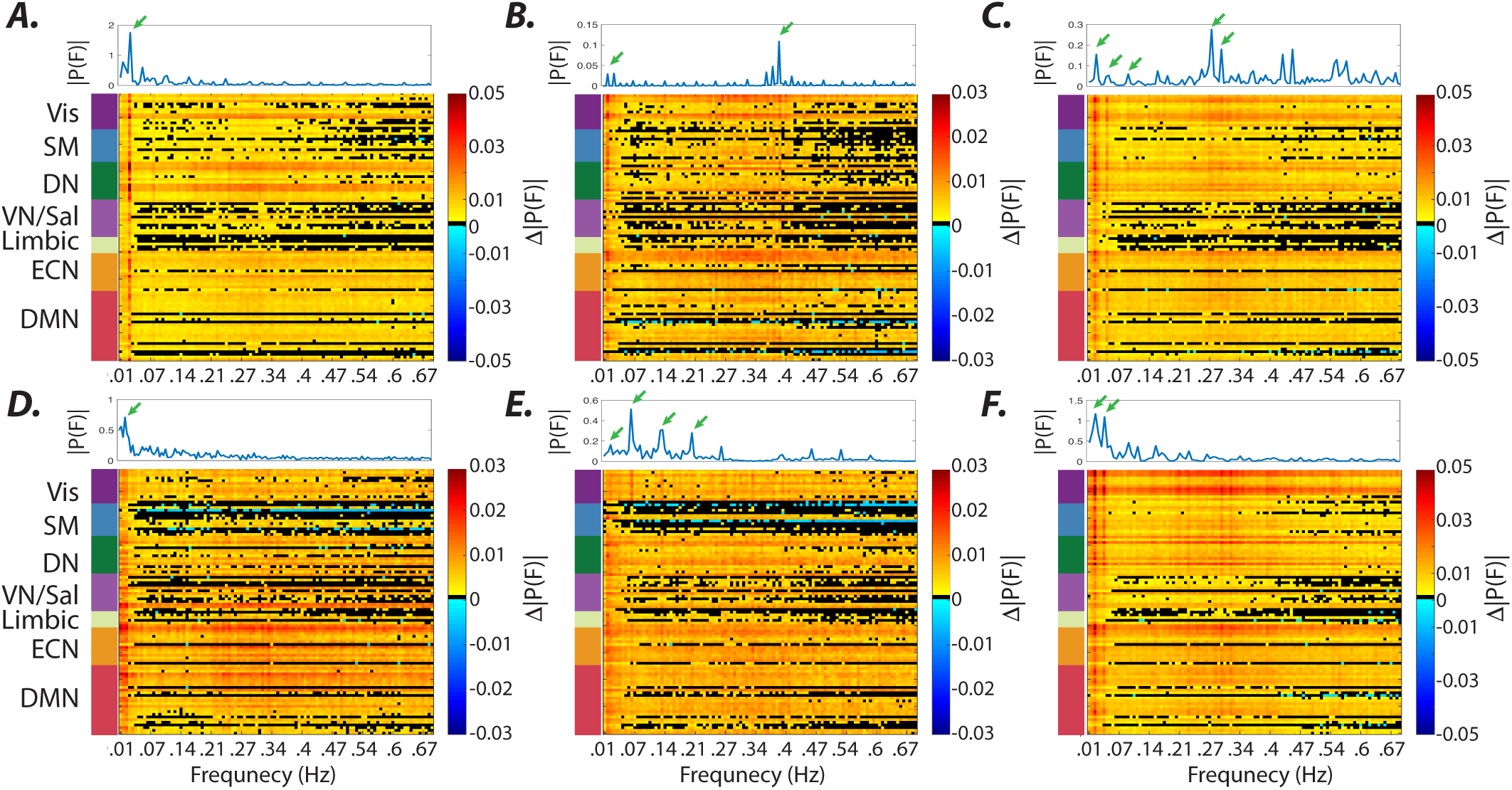
Matching spectral profiles of the known and estimated inputs. *(A-F)* The difference between average Fourier transform of the estimated inputs during task conditions (social, working memory, gambling, language, motor, and relational tasks, respectively) and resting state. Frequencies for which brain regions did not pass the significance level (Wilcoxon rank sum test, FDR *p* < 0.01) are represented in black. Top panels display the average (two sessions) spectral profile of the known boxcar regressors for each task (see SI-Fig. 3). Note that the expected frequency peaks of the external inputs (green arrows) are clearly identifiable across a wide rage of frequencies (even higher than 0.1 Hz). The brain regions are sorted and color-coded (left panel) based on the 7 resting state networks identified in (109), namely the visual (Vis), sensory/motor (SM), dorsal attention (DN), ventral attention/salience (VN/Sal), limbic, executive control (ECN), and default mode network (DMN).

**Figure 6.**
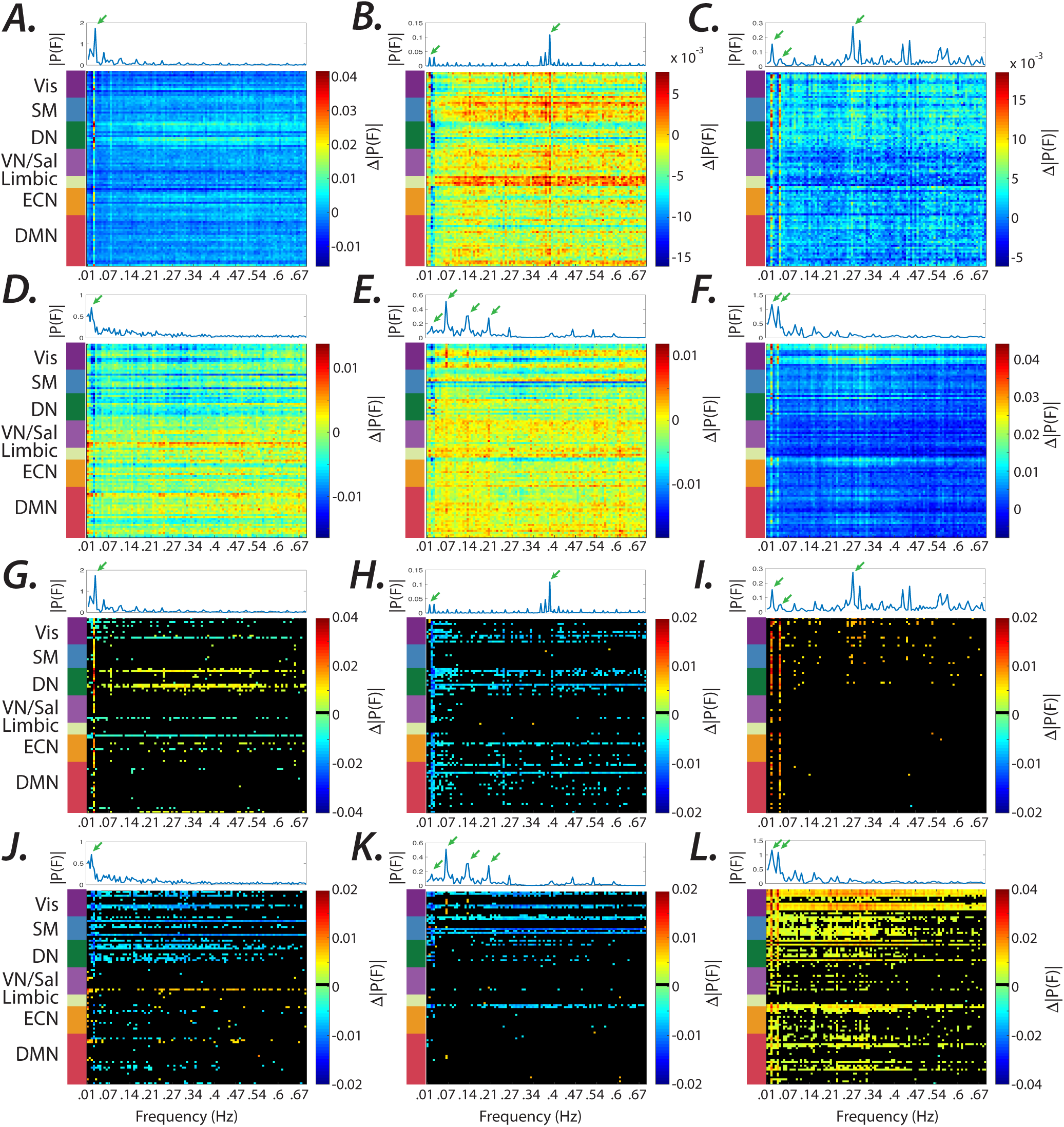
Task-related changes in the spectral profile of estimated inputs. *(A-F)* The difference between average Fourier transform of the estimated inputs during each task conditions (social, working memory, gambling, language, motor, and relational tasks) versus all other task conditions. Brain regions are sorted and color-coded (left panel) based on the 7 resting state networks identified in (109). Top panels displays the average (two sessions) spectral profiles of the known boxcar regressors for each task (see SI-Fig. 3). Note that the frequency peaks associated with different task conditions (marked by green arrow) are clearly echoed in the estimated inputs’ spectral power. ***(G-L)*** Non-parametric statistical testing (Wilcoxon rank sum, false discovery rate (FDR) *p* < 0.01) reveal the significance of these observations across several brain regions for all tasks over a wide range of frequencies. Brain regions did not pass the significance level are represented in black. Note that significant task-specific and spatially inhomogeneous changes in the estimated input’s spectral power can be found even as high as ≈ 0.2–0.3 Hz (e.g., panel ***(I)***).

**Figure 7.**
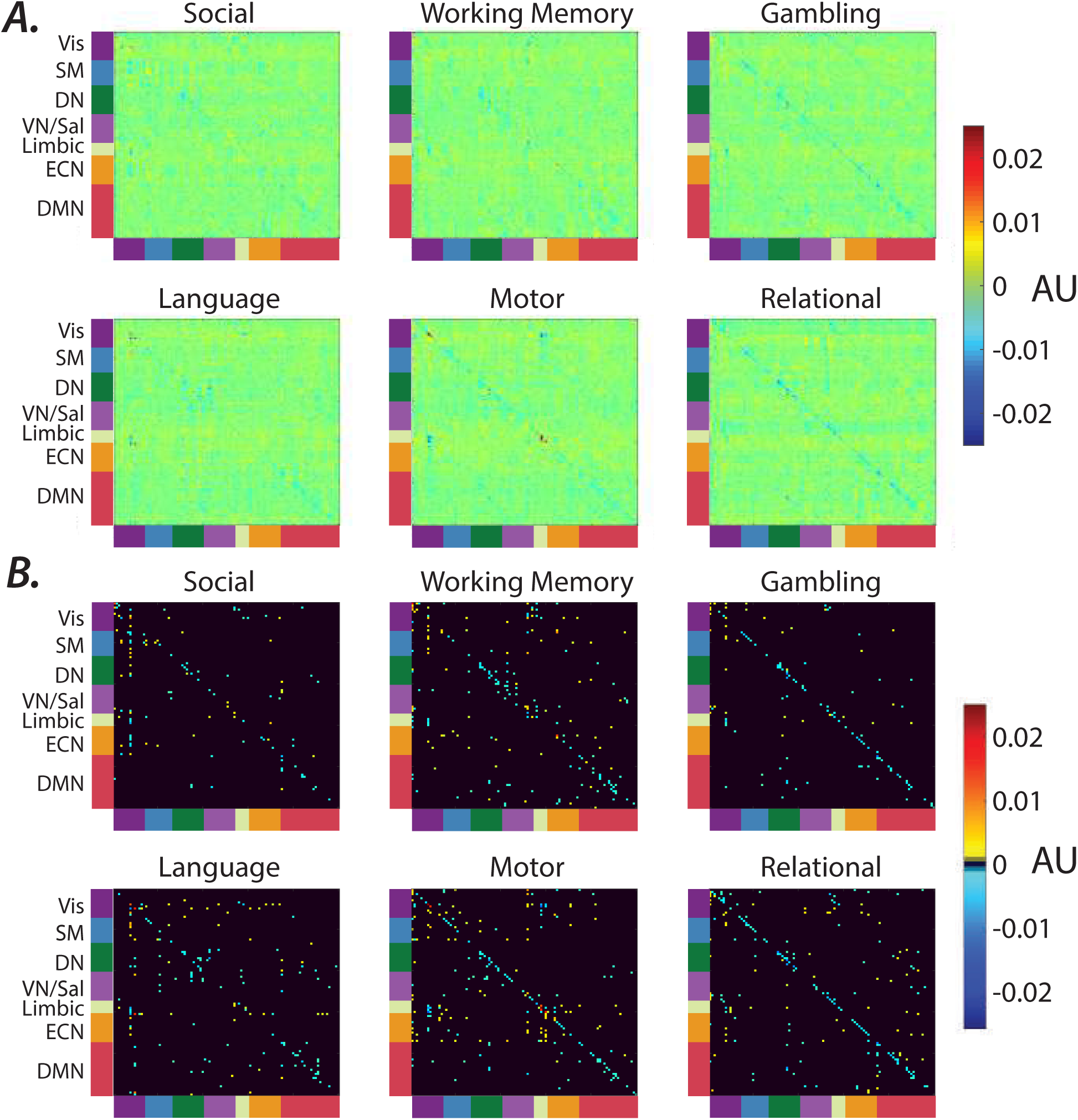
Task-related changes in the estimated system parameters. *(A)* Differences between the average identified system parameter (*A* matrix, for details see Materials and Methods) using a 210 TR window in task conditions versus resting state. The comparisons reveal task-related changes in the identified system parameters across all task conditions. ***(B)*** Same figures as in panel ***A***, except the parameters that did not passed the significance level are presented in black (Wilcoxon rank sum, false discovery rate (FDR) *p* < 0.01). Task-related changes are roughly characterized as reduced estimated self-excitation of large number of regions, while several regions known to be involved in the different tasks display elevated self-excitation, as well as altered interaction between different brain regions.

**Figure 8.**
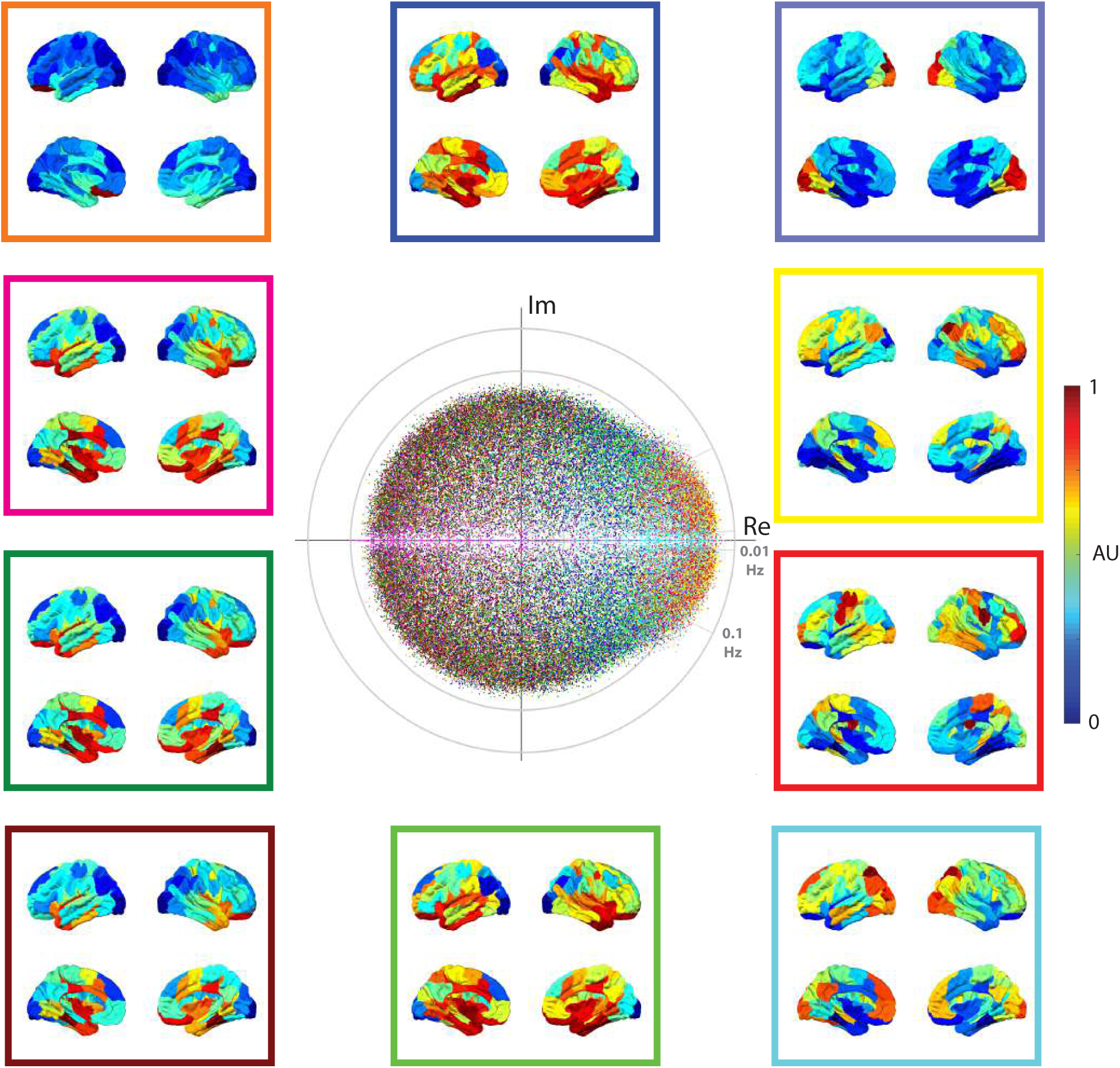
Distribution of eigenvalues estimated from the windowed (210 TR ≈ 2.5 mins) task time series. Here, the absolute of all eigenvectors estimated during task conditions are normalized and clustered into 10 clusters using *k*-means clustering algorithm. We color-code the clusters identified across all subjects, sessions, and windows (99 subjects × 4 sessions × 5 windows × 100 eigenmodes = 198,000 eigenvalues). The brain overlays represent the spatial distributions of eigenvectors associated with an eigenvalue (displayed with same color code) that is at the centroid of each cluster. To aid the visualization, the centroids are normalized after subtracting each centroid by its minimum element. The color bar represents the values of the normalized centroids. Although the estimated clusters during task are similar to that of the windowed resting state (Fig. 9), there are significant task-related changes present as described in SI-Fig. 11.

**Figure 9.**
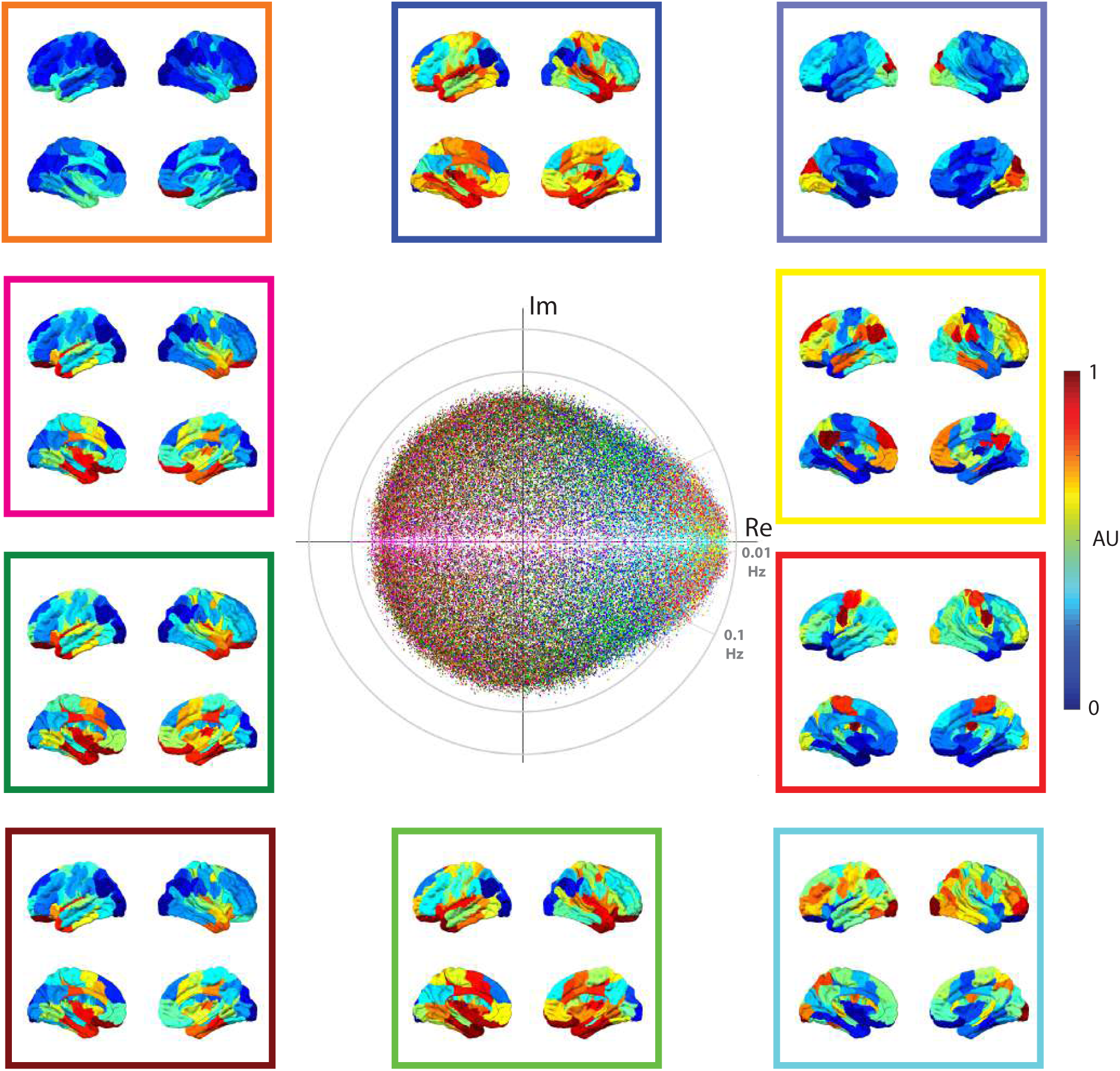
Distribution of eigenvalues estimated from the windowed (210 TR ≈ 2.5 mins) resting state time series. Here, the absolute of all eigenvectors estimated using short sliding window during resting state are normalized and clustered into 10 clusters using *k*-means clustering algorithm. Here, we color-code the clusters (*k* = 10) identified across all subjects, sessions, and windows (99 subjects × 4 sessions × 5 windows × 100 eigenmodes = 198,000 eigenvalues). The brain overlays represent the spatial distribution of eigenvector associated with an eigenvalue (displayed with same color code) that is at the centroid of each cluster. To aid the visualization, the centroids are normalized after subtracting each centroid by its minimum element. The color bar represents the values of the normalized centroids. Clustering (*k*-means) the eigenvalues based on their eigenvector’s similarity at shorter time periods reveals analogous to those shown in Fig. 2.

**Figure 10.**
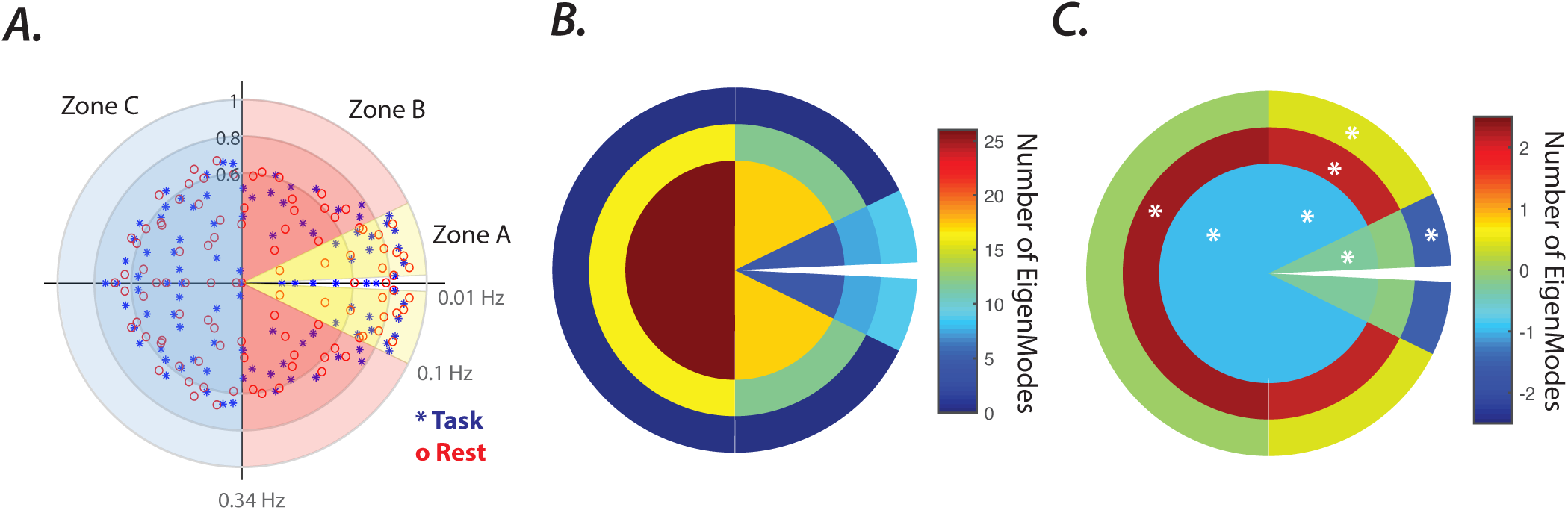
Task-related changes in the spatial profiles of eigenvector clusters. *(A-J)* The brain overlays represent the centroid of the 10 eigenvector clusters identified during windowed (210 TR ≈ 2.5 mins) resting state (SI-Fig. 9) and, task (SI-Fig. 8), as well as the normalized task-related changes in the cluster centroids (i.e., task-resting state), separately for each cluster. To aid the visualization, the centroids for both task and rest conditions were normalized after subtracting each centroid by its minimum element. Color bars represent the values of the normalized centroids. The task and resting state clusters were paired and color-coded similar to SI-Fig. 8 and SI-Fig. 9, based on the similarity (Pearson correlation) between cluster centroids during resting state and task as seen in SI-Fig. 13A. The high correlation coefficient (*r*) reveal the high overall similarity between the spatial profiles of the clusters between rest and task. Nevertheless, statistical comparison (Welch’s *t*-test, *p* < 0.05 FDR corrected for multiple comparisons across brain regions) highlight the brain-wide and significant task-related changes in cluster centroids. Brain regions that did not pass the significance-level are colored white. Interestingly, spatial profiles of task-related changes share similarities across clusters, as seen in SI-Fig. 13B. Furthermore, spatial profiles of the task-related changes also overlap with the ECN and DMN ROIs as seen SI-Fig. 13.

**Figure 11.**
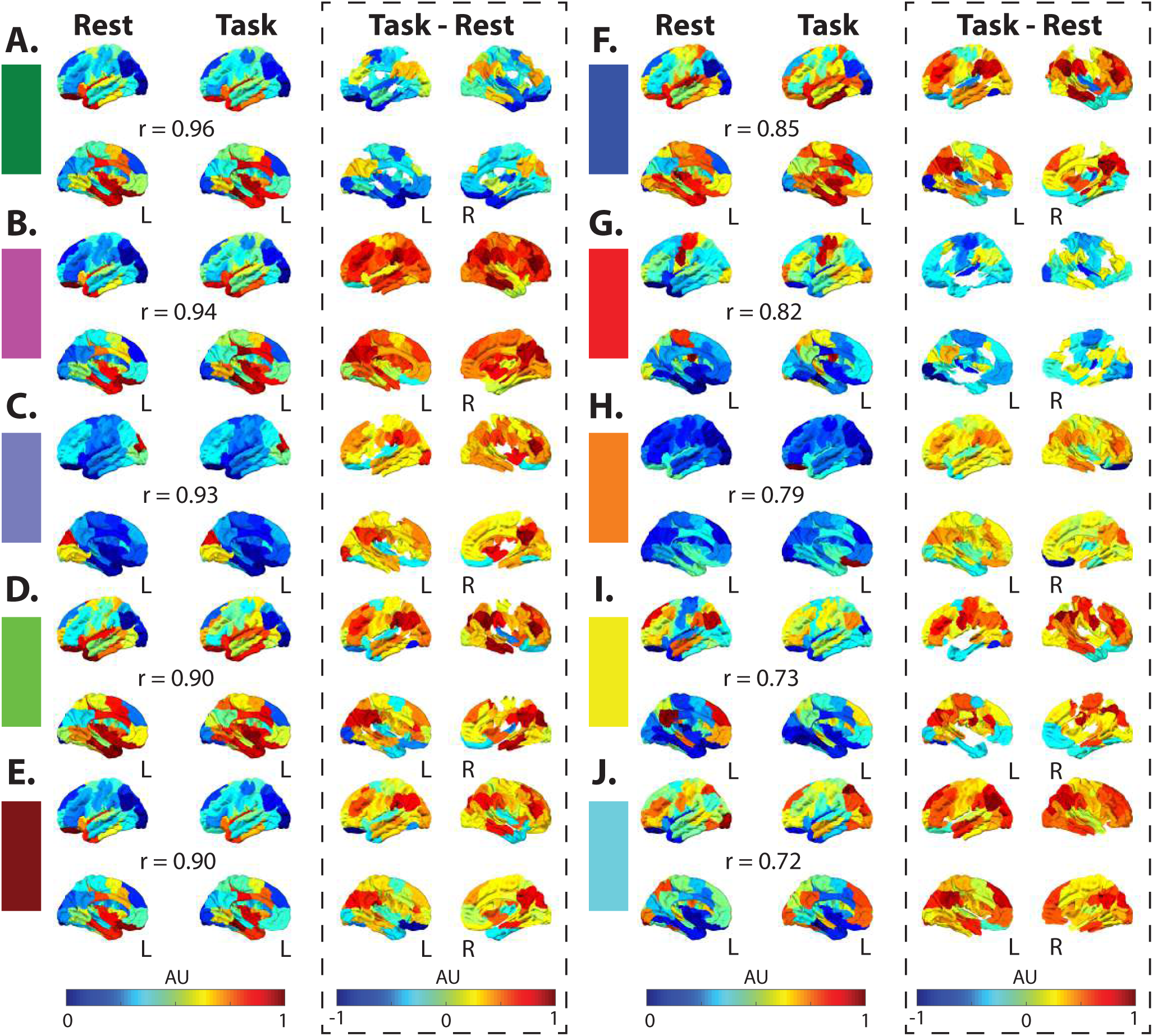
Task-related changes in the spatial profiles of eigenvector clusters. *(A-J)* The brain overlays represent the centroid of the 10 eigenvector clusters identified during windowed (210 TR ≈ 2.5 mins) resting state (SI-Fig. 9) and, task (SI-Fig. 8), as well as the normalized task-related changes in the cluster centroids (i.e., task - resting state), separately for each cluster. To aid the visualization, the centroids for both task and rest conditions were normalized after subtracting each centroid by its minimum element. Color bars represent the values of the normalized centroids. The task and resting state clusters were paired and color-coded similar to SI-Fig. 8 and SI-Fig. 9, based on the similarity (Pearson correlation) between cluster centroids during resting state and task as seen in SI-Fig. 13A. The high correlation coefficient (*r*) reveal the high overall similarity between the spatial profiles of the clusters between rest and task. Nevertheless, statistical comparison (Welch’s *t*-test, *p* < 0.05 FDR corrected for multiple comparisons across brain regions) highlight the brain-wide and significant task-related changes in cluster centroids. Brain regions that did not pass the significance-level are colored white. Interestingly, spatial profiles of task-related changes share similarities across clusters, as seen in SI-Fig. 13B. Furthermore, spatial profiles of the task-related changes also overlap with the ECN and DMN ROIs as seen SI-Fig. 13.

**Figure 12.**
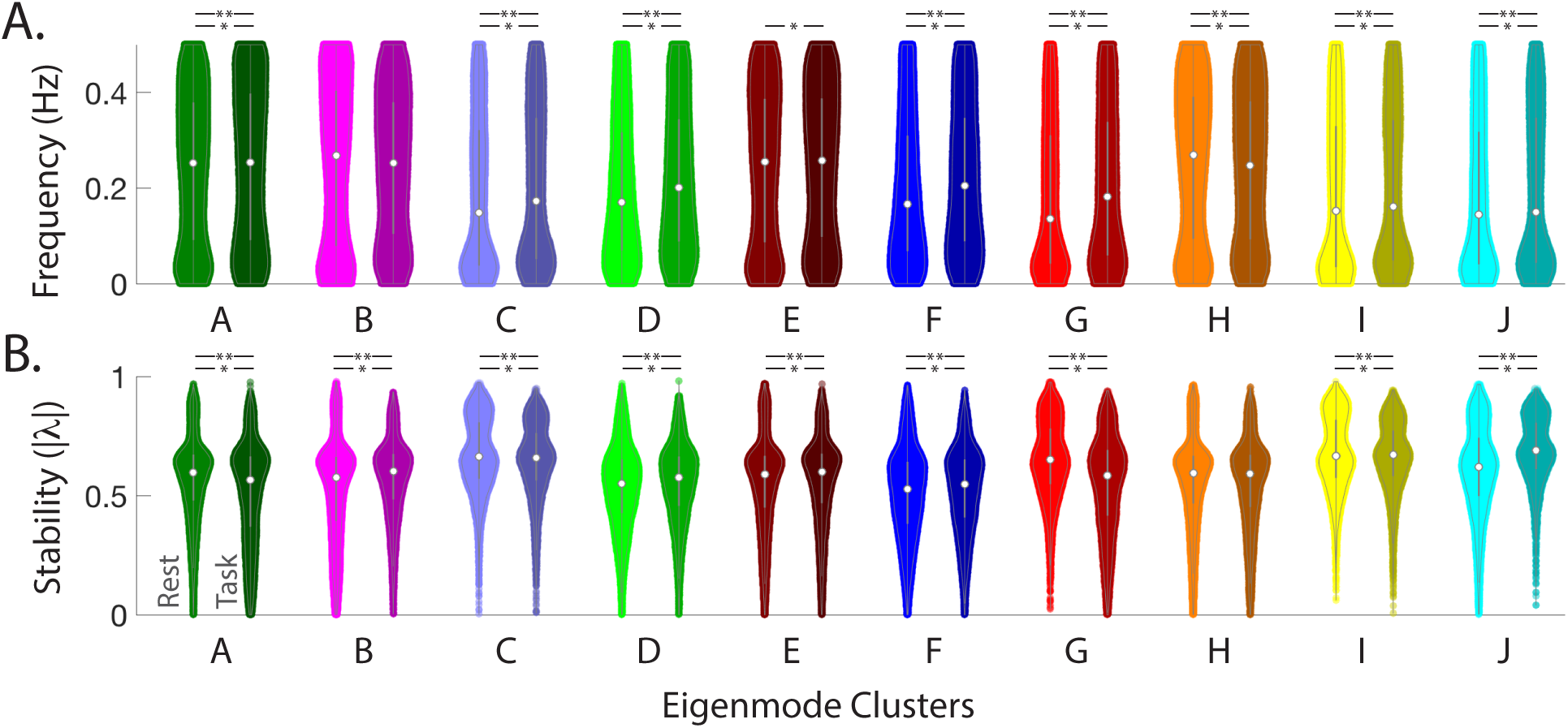
Task-related changes in the frequency and damping rate of eigenvector clusters. *(A)* Violin plots represent distributions of frequencies calculated from the eigenvalues associated with the eigenvector clusters, identified using 210 TR window during resting state (SI-Fig. 9), and during task (SI-Fig. 8). Plots are color-coded similar to SI-Fig. 11 and sorted in descending ordered from left to right, based on the similarity of cluster centroids during resting state and task (as seen in SI-Fig. 13A). Statistical comparisons between distributions of frequencies during resting state (left) and task (right, darker shade) using Wilcoxon rank sum test reveal significant task-related changes in most clusters (p < 0.001 marked by ‘*’, p < 0.0001 marked by ‘**’). ***(B)*** Distributions of eigenmode stability values (i.e., damping rate) calculated from the eigenvalues associated with each eigenvector clusters identified during resting state and task. Statistical comparisons between distributions of stabilities of eigenmodes (i.e., |*λ* |) during resting state (left) and task (right, darker shade) using Wilcoxon rank sum test also reveal significant task-related changes in most clusters (p < 0.001 marked by ‘*’, p < 0.0001 marked by ‘**’).

**Figure 13.**
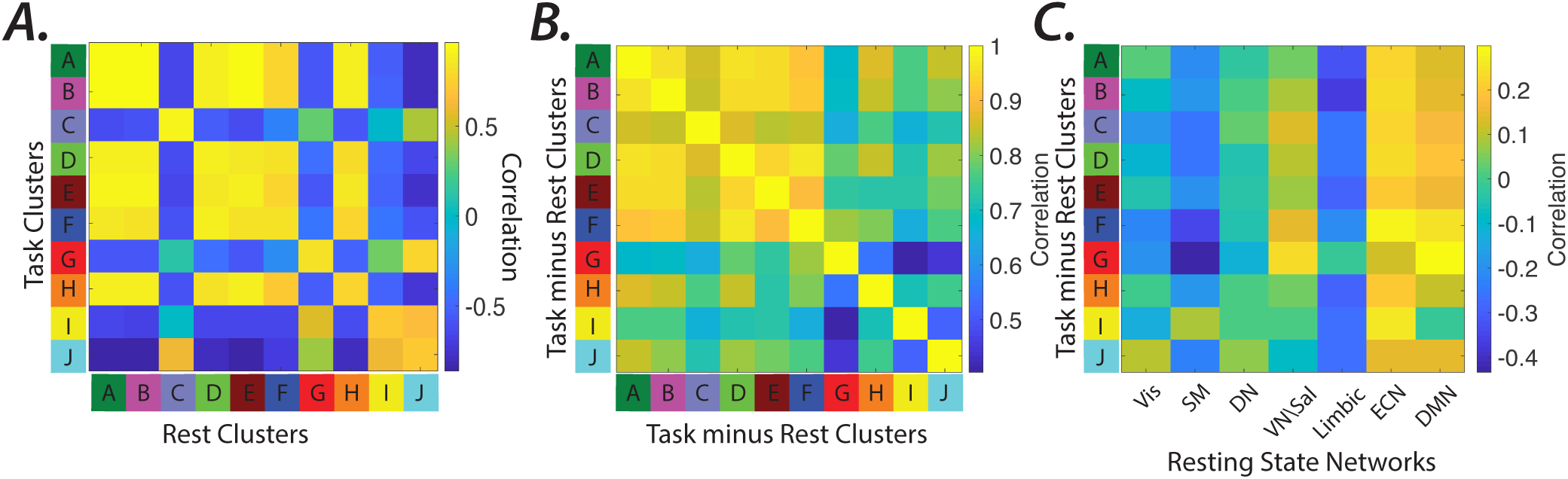
Similarity between eigenvector clusters identified during windowed resting state and task *(A)* The similarity matrix between cluster centroids during resting state and task. The values represent the Pearson correlation between the centroids of the clusters identified using a 210 TR window during task (SI-Fig. 8) and resting state (SI-Fig. 9). Cluster pairs with highest similarities were paired and ordered from the highest to lowest values (marked by A-J letters, respectively). The paired resting state and task clusters are color-coded similar to SI-Fig. 8 and SI-Fig. 9. ***(B)*** The similarity between the task-related changes in the cluster centroids. The matrix represents the correlation values between the task-related changes in the centroids of all possible cluster pairs. ***(C)*** The similarity between the task-related changes in the cluster centroids and resting state networks. The matrix represents the correlation values between the task-related changes in the cluster centroids and the 7 resting state network identified in (109).

**Figure 14.**
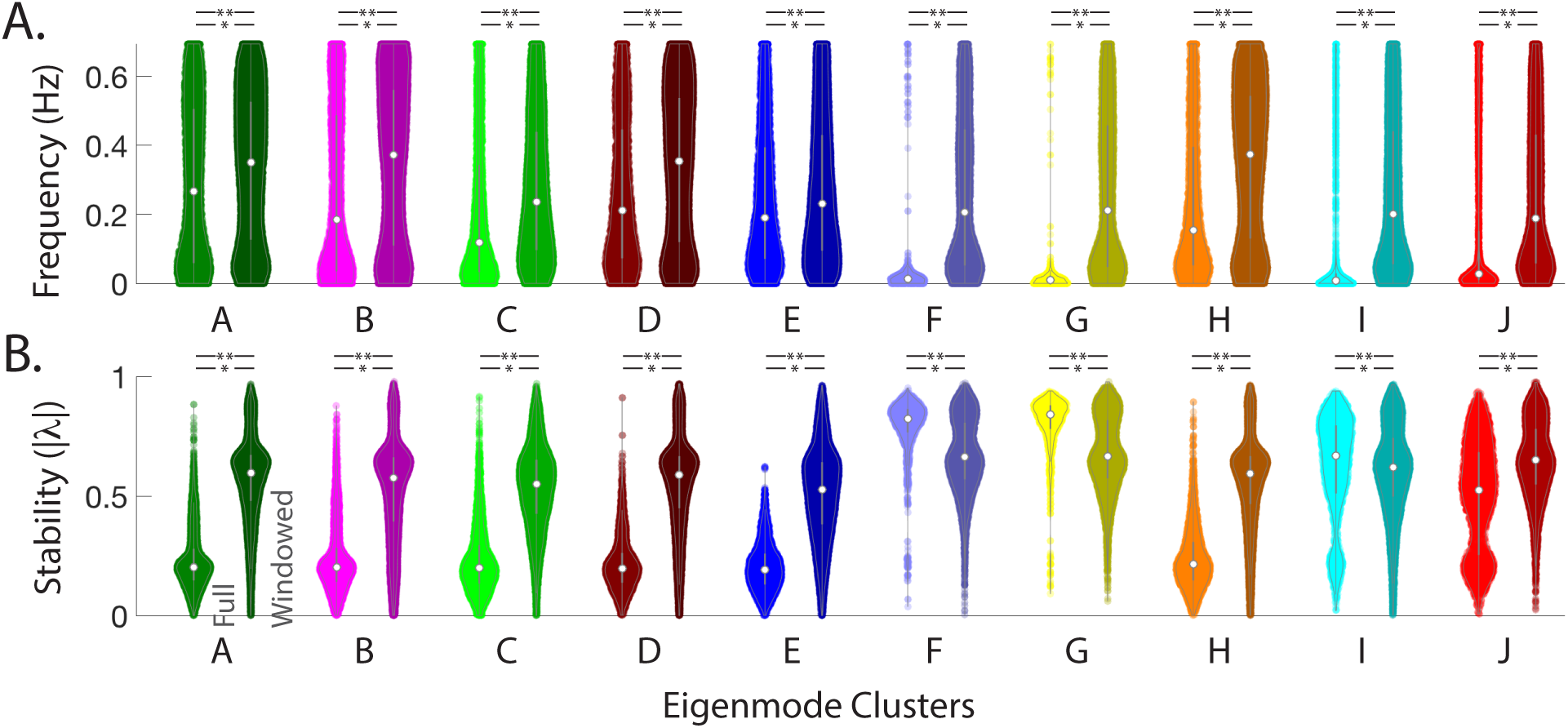
Differences in the frequency and damping rate of eigenvector clusters between full length and windowed resting state. *(A)* The violin plots represent distributions of frequencies calculated from the eigenvalues associated with the eigenvector clusters, identified during full length (Fig. 2), and windowed (210 TR) resting state (SI-Fig. 9). Plots are color-coded similar to SI-Fig. 2 and sorted in descending ordered from left to right, based on the similarity of the centroids of the clusters during resting state and task (as seen in SI-Fig. 13A). Statistical comparisons between distributions of frequencies during full length resting state (left) and windowed (right, darker shade) using Wilcoxon rank sum test reveal significant task-related changes in all clusters (p < 0.001 marked by ‘*’, p < 0.0001 marked by ‘**’). ***(B)*** Distributions of eigenmode stabilities (i.e., damping) calculated from the eigenvalues associated with the eigenvector clusters identified during windowed full length (left) and windowed (right, darker shade) resting state. Statistical comparisons between distributions of eigenmodes’ stability (i.e., *λ*) using Wilcoxon rank sum test also reveal significant differences in all clusters (p < 0.001 marked by ‘*’, p < 0.0001 marked by ‘**’) between the full length and windowed estimation at rest.

**Figure 15.**
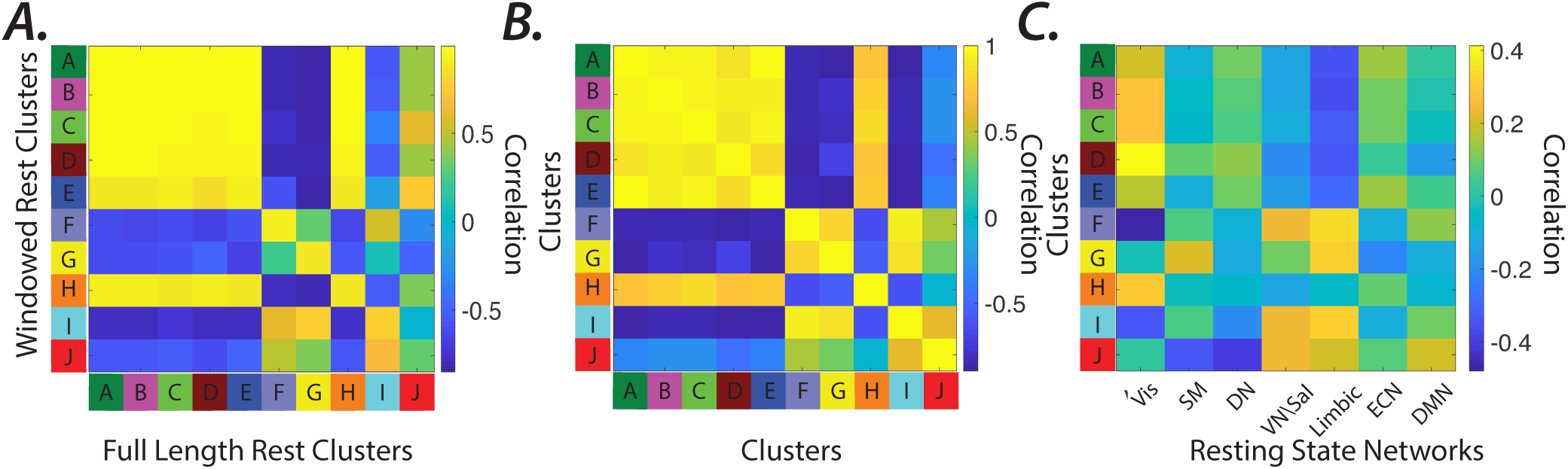
Similarity between eigenvector clusters identified during windowed and full length resting state *(A)* The similarity matrix between cluster centroids during windowed (210 TR ≈ 2.5 mins) and full length resting state (1200 TR ≈14.5 min). The values represent the Pearson correlation between the centroids of the clusters identified during full length (Fig. 2) and windowed resting state (SI-Fig. 9). Cluster pairs with highest similarities were matched from the highest to lowest values (marked by A-J letters, respectively). The paired windowed and full length resting state clusters are color-coded similar to Fig. 2. ***(B)*** The similarity (correlation) between the spatial profile of the difference between the full length and windowed resting state clusters. ***(C)*** The similarity between the difference between the full length and windowed resting state cluster centroids and the 7 resting state networks identified in (109).

**Figure 16.**
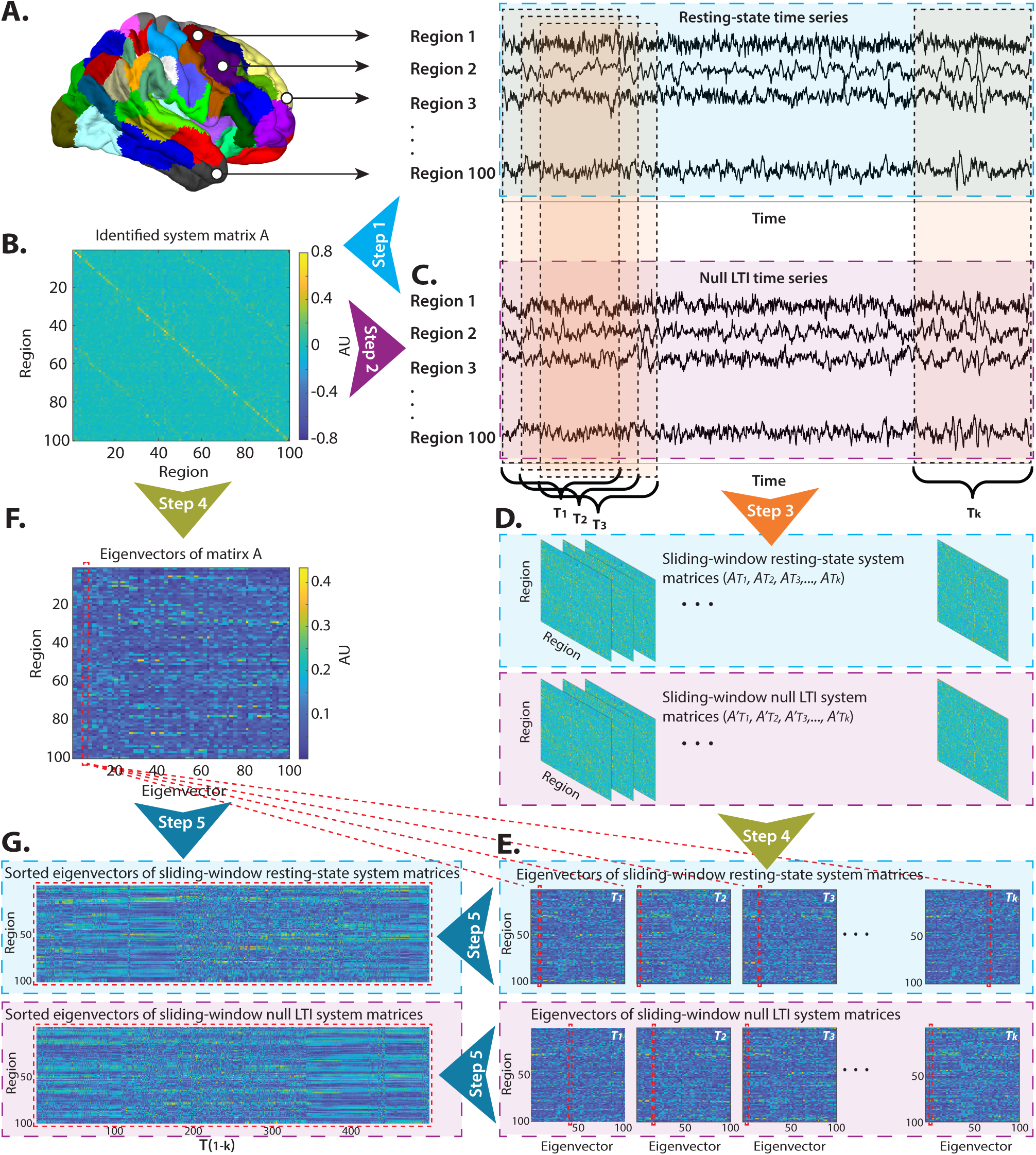
Testing the stationarity of the identified system during resting state scans. We hypothesize that system parameters are more accurately identified from the full length of the resting state scans (≈ 14.5 mins) and shorter windows introduce identification errors and lead to apparent system fluctuations. ***(A)*** To understand the extent of the expected fluctuations of a *true* LTI system, at first step, we identify all the parameters of an LTI system (the system matrix *A* in panel ***B***), the external input matrix *B*, the external inputs *U*, and the internal noise covariance matrix *e* from the full resting state time series (color-coded by cyan box). ***(C)*** Next, we simulate null LTI time series using these identified resting state parameters (color-coded by purple box). The residuals of model after fitting (at step 1) were added to the simulated time series, as a proxy for recording noise. ***(D)*** We hypothesize that the presence of non-stationarity inputs to the system result in apparent fluctuations in the identified systems. Furthermore, we conjecture that the extend of these fluctuations are higher in systems with true non-stationarities, and in systems with higher identification errors. Therefore, at step 3, we identified system matrices from both resting state and null LTI time series, using shorter sliding-window size of 210 TRs with 2 TR window increments (labeled as *T*_1_, *T*_2_, …, *T*_*k*_). The adjacency matrices in panel ***D*** represent the identified systems across *k*(= 496) increments of sliding identification windows. To identify which oscillatory modes of the system contribute the most to the observed non-stationarities, at step 4, we calculate the eigenvectors of all sliding-window system matrices (panel ***E***), as well as eigenvectors of matrix A (panel ***F***). Finally, at step 5, we sort all eigenvectors (according to their absolute value, | *V*_*A*_|) calculated from the sliding-window system matrices based on their similarity to the eigenvectors of the system identified from the full length resting state time series. See methods for more details on our sorting algorithm. ***(G)*** We provide examples of sorted eigenvectors from the sliding-window resting state and null LTI systems, based on a sample eigenvector of matrix A (highlighted in panel ***F*** with dashed red box). The matching eigenvectors across *k* window increments are also highlighted in panel ***E*** with dashed red box

**Figure 17.**
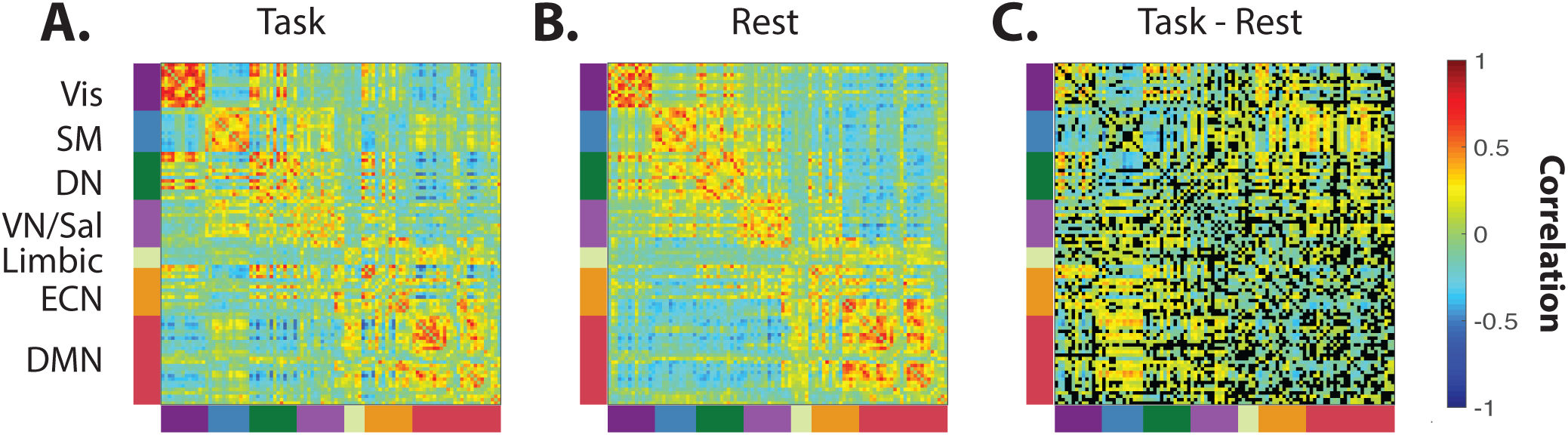
Task-related changes in the estimated inputs correlational structure *(A-B)* Average correlation matrix calculated from the estimated input during task and resting state scans, respectively. Brain regions are sorted and color-coded based on the 7 resting state networks identified in (109). ***(C)*** The task-related changes in the correlational structure of the estimated input is highlighted by subtracting **A** and **B** matrices, the pairs of brain region that did not show significant (*t*-test, FDR *p* < 0.01) difference in their correlation are colored black.

**Figure 18.**
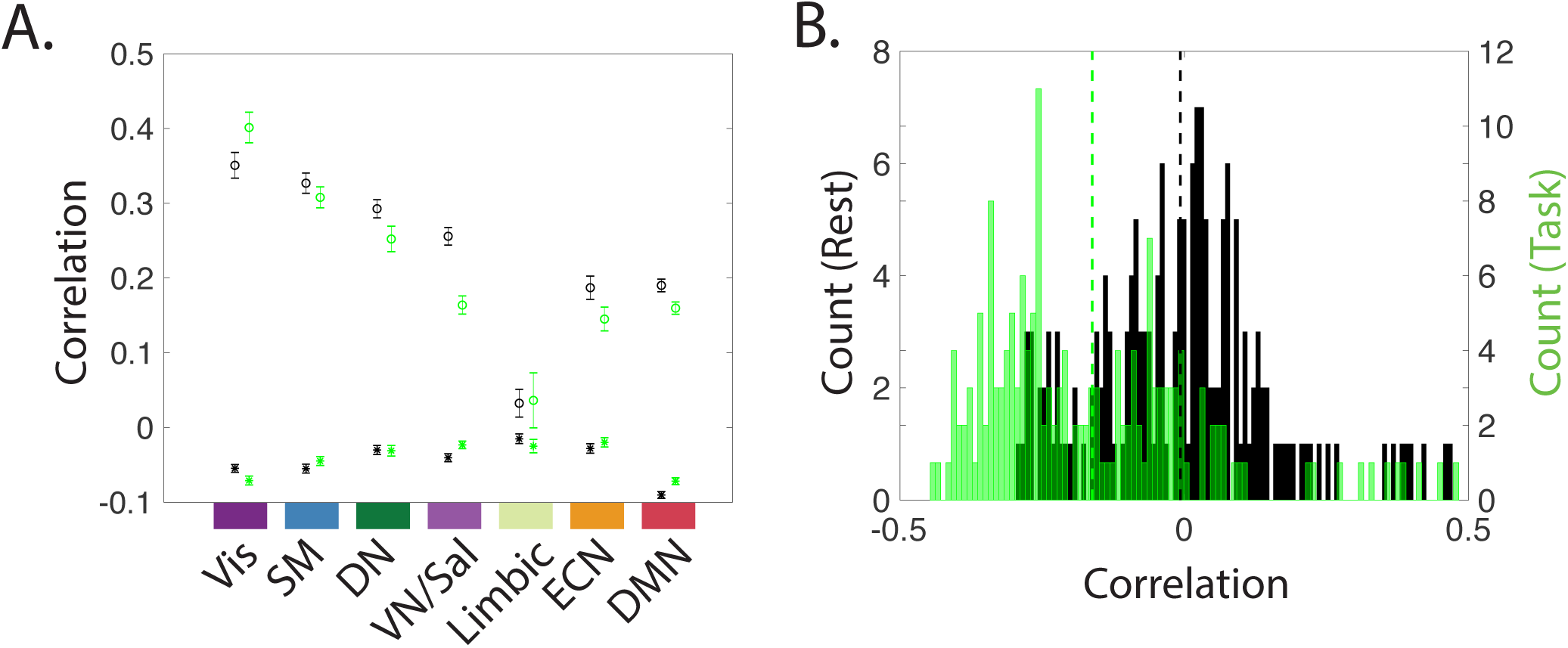
Within versus between resting state networks’ inputs correlation. *(A)* The average correlation between the estimated inputs to ROIs within 7 resting state networks (RSNs) identified in (109) during resting state (marked by red ‘o’) and during task (marked by blue ‘o’). We also presented the average correlation between the estimated inputs to to ROIs within and outside RSNs during resting state (marked by black ‘*’) and during task (marked by green ‘*’). The error bars represent the standard error. Statistical comparison (Wilcoxon rank sum, p < 0.001) reveals that, with the exception of Limbic network (resting state p = 0.0153, task p=0.0593), all RSNs display significantly higher within network correlations during both resting state and task. ***(B)*** Distributions of average correlation between SM and Vis ROIs during resting state (black), and during task (green). Statistical comparison between distributions reveals that the average correlation shifts significantly from low values at rest (r= −0.006±0.157, marked by vertical black line) to significantly (Wilcoxon rank sum, p < 0.001, p=3 ×10−17) higher anti-correlation values (r=-0.162±0.192, marked by vertical green line).

**Figure 19.**
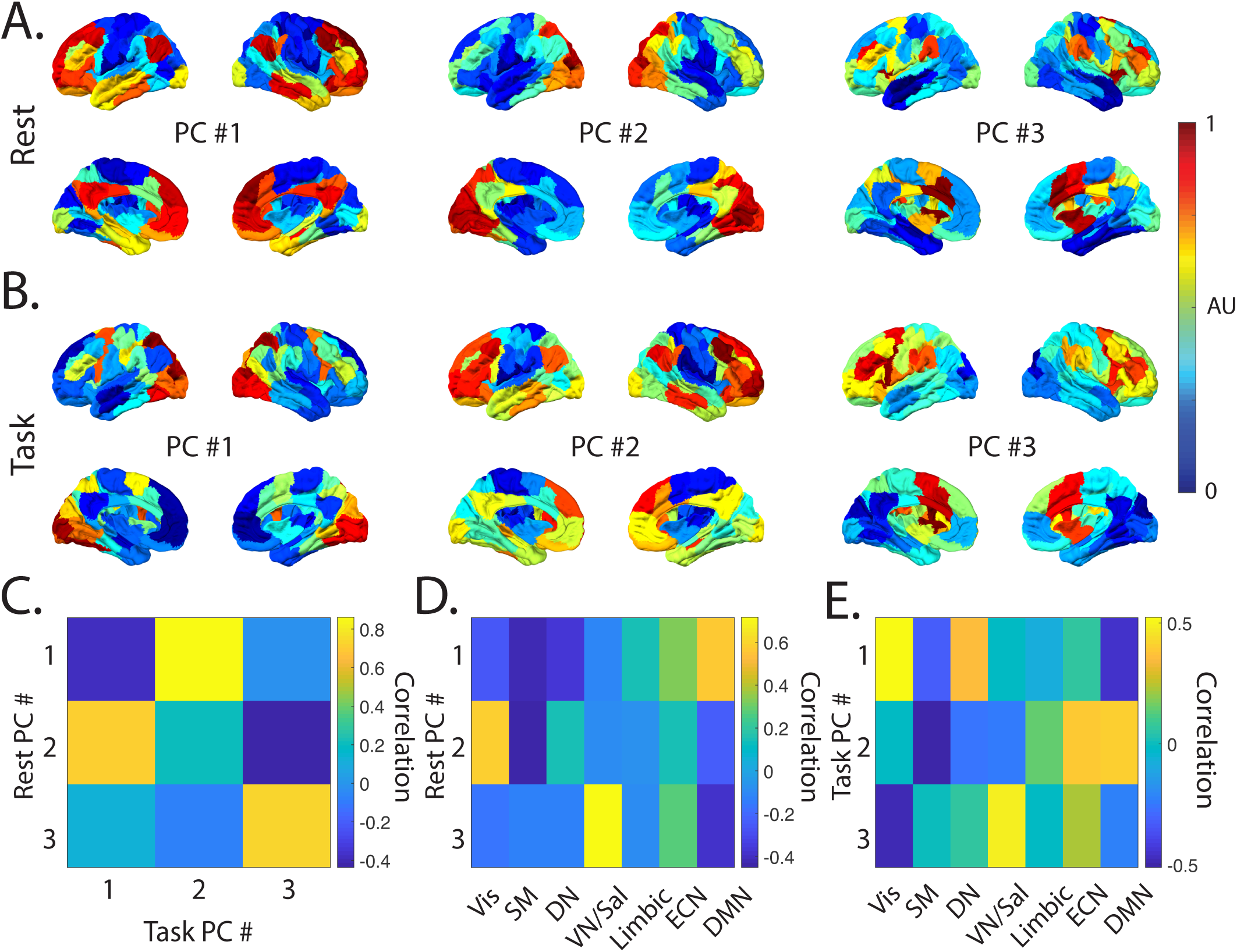
Principle components of the system’s input matrices B *(A*–*B)* The brain overlays display the top three principal components (sorted based on explained variance) calculated from the concatenated B matrices (all subjects) during resting state and task scans, respectively. The color bar represents the values of the normalized principal components. ***(C)*** Similarity matrix represents the correlation values between three principal components presented in panels ***A*** and ***B. (D*–*E)*** Similarity matrix represents the correlation values between three principal components presented in panels ***A*** and ***B*** and the 7 resting state networks identified in (109), respectively.

**Figure 20.**
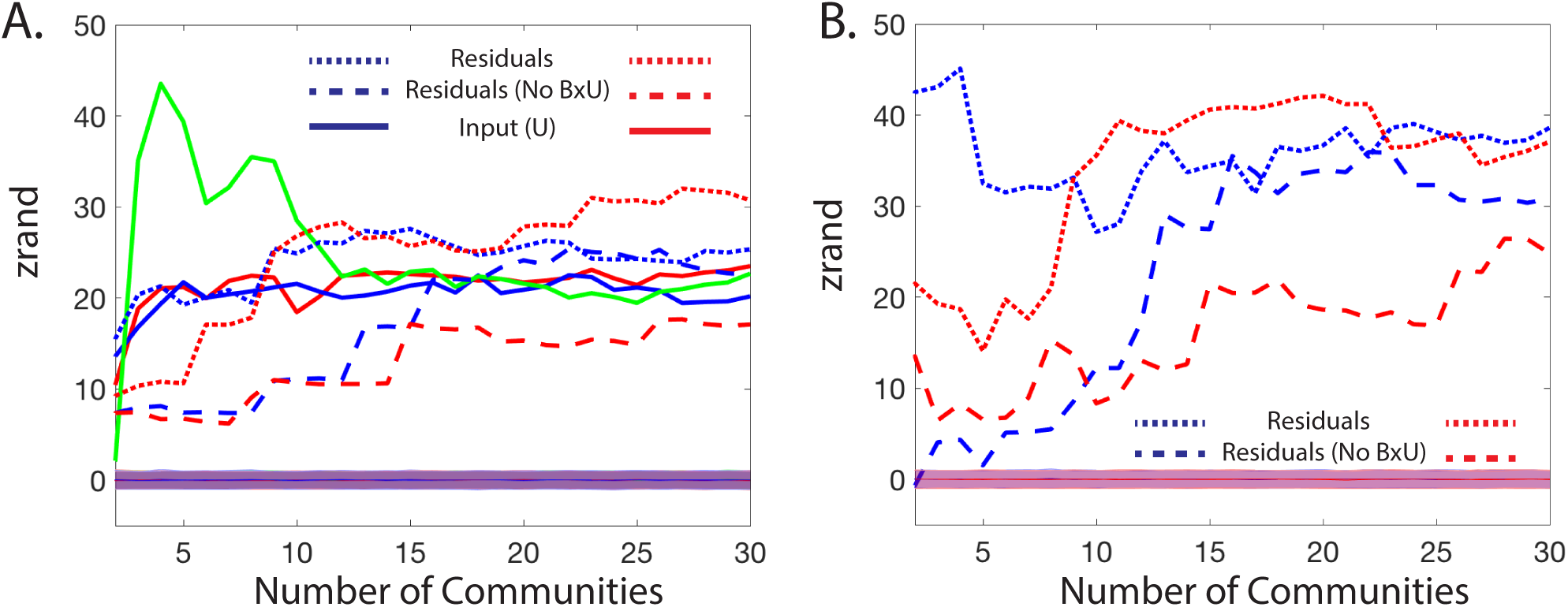
The correlational structure of estimated inputs and model residuals. *(A)* The average similarity, measured by the z-score of the Rand coefficient, between the hierarchical community organization of the correlational structure of the model’s residual and inputs to 17 resting state networks identified in (109) estimated during resting state (blue) and task conditions (rest). The shaded curves show the average similarity and standard deviation of the nulls, calculated by randomizing the community labels (*n* =50,000). The green line represents the similarity between communities identified in the task and resting state at different topological scales (i.e., number of communities). ***(B)*** The average z-score of the Rand coefficient (112) between the hierarchical community organization of the correlational structure of the model’s residual to the estimated inputs, during resting state (blue) and task conditions (red). The shaded curves show the average similarity and standard deviation of the nulls, calculated by randomizing the community labels (*n* =50,000).

